# VEXAS anemia is a mosaic erythroblastopenia

**DOI:** 10.1101/2024.12.02.623560

**Authors:** François Rodrigues, Giulia Hardouin, Sara El Hoss, Aya Ghoul, Emilie-Fleur Gautier, Michaël Dussiot, Sandy Peltier, Pascal Amireault, Vanessa Soldan, Annarita Miccio, Mounia Debili, Vincent Jachiet, Thiago Trovati Maciel, Julien Rossignol, Eric Allemand, Arsène Mekinian, Sophie Georgin-Lavialle, Mohammad Salma, Eric Soler, Pierre-Emmanuel Gleizes, Marie-Françoise O’Donohue, Olivier Kosmider, Manuel S. Rodriguez, Olivier Hermine, French VEXAS group, MINHEMON

## Abstract

VEXAS (vacuoles, E1 enzyme, X-linked, autoinflammatory, somatic) is a recently discovered autoinflammatory disorder linked to somatic mutations in the *UBA1* gene, resulting in a profound cytoplasm-restricted defect in ubiquitylation. The disease is characterized by a macrocytic anemia that remains poorly understood. To investigate the erythroid lineage in VEXAS, we conducted a comprehensive study combining *in vivo* assessments of patients’ mature red cells and marrow erythroblasts, alongside *in vitro* base-editing models of erythropoiesis. Here we show that mature red cells do not exhibit ubiquitylation defects, and patient-derived bone marrow erythroblasts lack *UBA1* mutations beyond the basophilic stage of erythroid differentiation. *In vitro* base editing of *UBA1* variants in CD34+ primary cells resulted in high mortality during early erythroid differentiation, but not during monocytic differentiation. Edited erythroid precursors displayed TP53 overexpression linked to defective ubiquitylation and anomalies in ribosome biogenesis, reminiscent of Diamond-Blackfan anemia. We propose that VEXAS-associated anemia should be considered as a mosaic erythroblastopenia, where the severity of anemia is influenced by the quality and quantity of the *UBA1-*WT compartment. These insights may aid clinicians in tailoring treatment strategies.

## Introduction

VEXAS (vacuoles, E1 enzyme, X-linked, autoinflammatory, somatic) is a recently identified autoinflammatory disorder first described by Beck et al. in 2020^1^. It is caused by somatic missense mutations in hematopoietic stem cells (HPSCs), specifically affecting methionine-41 (p.Met41) or splice sites in the *UBA1* gene located on the X chromosome^1^. The E1 ubiquitin-activating enzyme UBA1, which is ubiquitously expressed, plays a crucial role in the activation of ubiquitin, marking the initial step of the ubiquitylation cascade^2^. To date, only two enzymes responsible for the E1 step of ubiquitylation have been identified in humans: UBA1 and UBA6, the latter exhibiting a significantly lower expression level in tissues^3,4^.

Mutations leading to VEXAS disrupt translation of the canonical cytoplasmic isoform of UBA1 (UBA1b), resulting in its replacement by a truncated, catalytically impaired cytoplasmic isoform (UBA1c), and causing a profound defect in cytoplasmic ubiquitylation^1^. Notably, these mutations do not affect the function of the nuclear isoform of UBA1 (UBA1a). Clinically, VEXAS manifests through a spectrum of symptoms, including fever, skin lesions, pulmonary inflammation, chondritis of the ears and nose, and both arterial and venous inflammation and thrombosis^1,3^. Hematologic abnormalities include macrocytic anemia that can be profound, thrombocytopenia, and monocytopenia. However, anemia is in the majority of cases mild and not requiring transfusions^4^.

Current therapeutic strategies for VEXAS are not well-defined and primarily focus on reducing inflammation and managing anemia. In many VEXAS patients, the disease presents as a mosaic of mutated (mut-*UBA1*) and wild-type (wt-*UBA1*) cells, with additional mutations associated with clonal hematopoiesis and myelodysplastic syndromes observed in the bone marrow. While somatic mutations at p.Met41 have been identified in hematopoietic stem cells and myeloid cells in peripheral blood, they are absent in B and T cells^1^.

To date, there has been no dedicated investigation into erythropoiesis and red blood cells (RBC) characteristics in VEXAS patients. It remains unclear whether circulating RBC in these patients originate from mut-*UBA1* or wt-*UBA1* bone marrow clones. If erythropoiesis can be achieved despite *UBA1* mutations, mut-*UBA1* RBCs presenting abnormal proteomes could potentially contribute to peripheral inflammation, similar to conditions such as sickle cell disease, lupus, or CANDLE syndrome, where RBC anomalies provoke inflammation through the release of stressors during hemophagocytosis or hemolysis^5,6^. Conversely, if mut-*UBA1* erythropoiesis is inefficient or defective, this would provide new insights into the role of UBA1 and proteostasis during erythropoiesis and into the treatment strategies for anemia in VEXAS.

To investigate the red cell lineage in VEXAS, we carried out an *in vivo* assessment of RBCs and marrow erythroblasts in patients, alongside *in vitro* base editing experiments introducing VEXAS-causing mutations into CD34+ cells cultured towards the erythroid lineage. Through this research, we aim to elucidate the impact of *UBA1* mutations on erythropoiesis and their potential role in the pathophysiology of VEXAS, ultimately contributing to improved treatment strategies for patients who continue to face significant unmet medical needs.

## Materials and methods

### Erythrocyte sample preparation

Approximately 3–8 mL of peripheral blood in EDTA tubes were obtained from patients and healthy individuals with informed written consent according to the Declaration of Helsinki (ethics approval number GR-Ex/CPP-DC2016-2618/CNIL-MR001). Whole blood was centrifugated 10 minutes at 450 g. Plasma and buffy coat were removed and aliquots of 2 uL of red cell pellet were washed with PBS and used for downstream flow cytometry assays. For preparation of nLC-MS/MS proteomic samples, 2 µL of pellet were collected per condition. After washing with PBS, the pellets were incubated with 1 µL of anti-GPA antibody (562938, BD Biosciences) for 15 minutes. After washing, cells were resuspended in thiazole orange and GPA^pos^ thiazol orange^neg^ cells were sorted with BD Aria II cell sorter. After sorting, cells were washed three times in PBS before being solubilized in lysis buffer.

### Ghost preparation

Ten million sorted erythrocytes per condition were incubated for 20 minutes at 4°C in hypotonic 5P8 buffer (5 mM Na_2_PO_4_, pH 8, 0.35 mM EDTA, 1 mM phenylmethylsulfonyl fluoride, reference 36978 Thermofisher). The membranes were pelleted by centrifugation for 15 minutes at 22 000*g*. The ghost pellet was washed several times with hypotonic buffer to obtain a white pellet which was solubilized in lysis buffer.

### Sample preparation for nLC-MS/MS proteomic analysis

Five million of sorted erythrocytes and 10 million of corresponding ghosts per sample were boiled in Tris HCl 200mM pH 8.5 SDS 2% lysis buffer for 5 min at 95°C. Protein concentrations were calculated in erythrocyte samples using a bicinchoninic acid assay (BCA kit; Pierce). Disulfide bridges from 50 µg of proteins for total erythrocyte samples and the totality of proteins for ghosts samples were then reduced using TCEP 20 mM and subsequent free thiols groups were protected using chloroacetamide 50 mM for 5 min at 95°C. Proteins were trypsin-digested overnight using filtered-aided-sample-preparation method as described previously (Wisniewski JR et al., Nat Meth, 2009). Eluted peptides were fractionated into 5 fractions using strong cation exchange (SCX) StageTips and vaccum-dried while centrifuged in a Speed Vac (Eppendorf).

### nLC-MS/MS proteomic analysis

Mass spectrometry (MS) analyses were performed on a Dionex U3000 RSLC nano-LC system coupled to an Orbitrap Fusion mass spectrometer (both from Thermo Fisher Scientific). Peptides from each SCX fraction were solubilized in 0.1% trifluoracetic acid containing 10% acetonitrile and loaded, concentrated and washed on a C18 reverse phase precolumn (3-µm particle size, 100 Å pore size, 75-µm inner diameter, 2-cm length; Thermo Fischer Scientific). A tenth of peptides from total erythrocyte digest and a quarter of peptides from ghosts digests were then separated an Aurora C18 reverse phase resin (1.6 μm particle size, 100Å pore size, 75μm inner diameter, 25cm length mounted to the Captive nanoSpray Ionisation module, from IonOpticks, Middle Camberwell Australia) with a 3-hour gradient starting from 99% solvent A (0.1% formic acid) and ending with 55 % solvent B (80 % acetonitrile, 0.085% formic acid). The mass spectrometer acquired data throughout the elution process and operated in a data-dependent scheme with full MS scans acquired with the Orbitrap, followed by as many MS/MS ion trap HCD spectra 2 seconds can fit (data-dependent acquisition with top speed mode: 2-s cycle) using the following settings for full MS: automatic gain control (AGC) target type: custom with a normalized AGC target of 50%, maximum ion injection time (MIIT): 60ms, isotopes exclusion, resolution: 6.10e4, m/z range 350–1500. For HCD MS/MS: Quadrupole filtering, Normalised Collision Energy: 30. Ion trap rapid detection: isolation width: 1.6 Th, minimum signal threshold: 5000, AGC: 2.10e 5, MIIT: 60 ms, resolution: 3.10^4^. A dynamic exclusion time was set at 30 s. Peptides with charge state under than 1 or upper than 7 were excluded from fragmentation.

### Data processing protocol

Identifications and quantifications of proteins were performed using MaxQuant version 1.6.17.0^7^ the reviewed Human Uniprot sequence database without isoforms (release Dec 2023) and a list of frequent contaminant sequences. The false discovery rate was kept below 1% on both peptides and proteins and a maximum 2 missed cleavages was allowed. Carbamidomethylation of cysteines was set as constant modification and acetylation of the protein N terminus and oxidation of methionine were set as variable modifications. Label free protein quantification (LFQ) was done using both unique and razor peptides with at least 2 ratio counts. The “match between runs” (MBR) option was allowed.

For statistical analysis, data were imported into the Perseus software version 1.6.14.0^8^. Reverse and contaminant and only identified by site proteins were excluded from analysis. Absolute quantification in copy number per cell for each protein for total erythrocyte proteomes was done using measured MCH values as previously described^9^. For ghost proteomes absolute quantifications, calculated BAND3 amount in the corresponding total proteome was used as a reference. Only proteins quantified in at least 3 samples of one condition were selected for two samples Student’s T-Test.

### Morphologic analysis of red cells using imaging flow cytometry

Imaging flow cytometry (ImageStream X Mark II, AMNIS part of Cytek Biosciences) was performed to determine RBC dimensions and morphology by using brightfield images (×60 magnification) processed with computer software (IDEAS v6.2, AMNIS) as previously reported^10^. Ten µL of whole blood where resuspended at a 1:100 dilution in a Krebs–albumin solution (Krebs–Henseleit buffer, Sigma-Aldrich) modified with 2.1 g of sodium bicarbonate, 0.175 g of calcium chloride dehydrate, and 5 g of lipid-rich bovine serum albumin (Albu-MAX II, Thermo Fisher Scientific) for 1 L of sterile water (pH 7.4). Focused RBCs and single RBCs were selected using the features Gradient RMS_M01_Ch01 and Aspect ratio_-M01_Ch01 vs Area_M01_Ch01. Finally, front views were selected using the feature Circularity_Object (M01, Ch01, Tight), and projected surface area was determined using the feature Area_Object (M01, Ch01, Tight). Mean projected surface area between groups was compared with a two-tailed Mann-Whitney test.

### RBC elongation index measurement

Elongation capacity of RBC was evaluated by measuring the elongation index (EI) by ektacytometry (LORRCA MaxSis, RRMechatronics) at various shear stresses of (0.3, 0.53, 0.95, 1.96, 5.33, 9.49, 16.87 and 30 Pa) as previously described^10^. Groups were compared using a two-way ANOVA test.

### Flow cytometry assays

Two uL of the red cell pellet was washed with PBS stained with lactadherin or thiazole orange, incubated for 15 minutes and analysed on a Beckman Coulter Gallios. For mitochondrial content in mature RBCs, 5 million red blood cells per condition were resuspended in RPMI with MitoTracker Deep Red (MTDR, 200 nM, Thermo-Fisher) +/- FCCP (Carbonyl cyanide-p-trifluoromethoxyphenylhydrazone, 500 nM) and incubated for 30 minutes at 37°C. The cell pellet was then washed four times with PBS, treated with Fc Block (BD Bioscience) and stained with human anti-235a and FITC anti-human CD71. The fluorescence intensity of MTDR was measured, in the mature CD71^neg^ CD235a^pos^ RBC population on a Beckman Coulter Gallios. For flow cytometry differentiation analyses, cells were washed with PBS at the indicated time and then stained for GPA/Band3/CD49d expression with 1:20 human BV421-anti-235a (Beckman Coulter), 1:20 human FITC-BRIC6 (IBGRL Research) and 1:20 human APC-anti-CD49d (Beckman Coulter). For viability analyses, cells were washed with PBS, resuspended in 1 X annexin buffer (Biolegend) and labeled with annexin V-APC (BD Biosciences) and 7-AAD or propidium iodide. FACS experiments were conducted on a Beckman Coulter Gallios or BD Fortessa Flow cytometer. The data were analyzed using FlowJo software (version 10.0.8, TreeStar) and significance of differences was evaluated by a two-tailed Mann-Whitney test.

### Cell sorting of patients’ bone marrow erythroblasts

Mononuclear cells were isolated from bone marrow by Ficoll centrifugation. After two PBS washes, cells were stained with anti-glycophorin, anti-Band3, anti-CD49d antibodies for 20 minutes at 4°C. After another PBS wash, they were sorted using the Aria II BD SORP sorter to isolate the following populations: non-erythroid (GPA^neg^), basophils and polychromatophils (Band3^pos^CD49d^pos^), orthochromatophils (Band3^pos^CD49d^neg^). DNA was isolated from frozen sorted pellets using the Qiagen DNA Micro Kit and the beginning of UBA1 exon 3 was amplified by PCR before Sanger sequencing.

### Primary cell culture

CD34 + cells were isolated from umbilical cord blood (CD34 beads, LS columns, Miltenyi) obtained from male donors (CRB Saint-Louis Hospital Paris, France) and mature erythroid cells were generated in a two-step amplification culture as previously described^11^, that is, with a first eight-day long EPO-free phase (phase I, figure 3A, SCF 100 ng/mL, IL-3 10 ng/mL), followed by a second phase with EPO (phase II, figure 3A, EPO 2 UI/mL, SCF 50 ng/L for seven days with IL-3 10 ng/mL the first three days, followed by EPO 3 UI/mL SCF 20 ng/L, FBS 3% until the end of the culture). In our notation system, day 8 of phase I = day 0 of phase II. The CD36^neg^ fraction of sorted cells at day 0 of phase II was cultured towards the myelomonocytic lineage as previously described^12^ with 50 ng/mL SCF and 25 ng/mL M-CSF, with the addition of 20 ng/mL IL-3.

Patient peripheral blood CD34+ cells were obtained after ethical approval and written informed consent (GR-Ex/CPP-DC2016-2618/CNIL-MR001) and cultured with another protocol to compensate for low initial cell numbers as well as aging and disease-related proliferation defects. That is, as previously described^13^ with the following modifications : addition of 10% FBS from day 0 to 7 and 3% FBS from day 7 to 15, addition of 10 ng/mL IL-6 from day 0 to 7, EPO concentration 3 UI/mL from day 0 to 15.

### HUDEP-2 culture

HUDEP-2 cells were cultured as previously described^14^ with the following modifications : in the amplification phase, Stemspan was replaced with IMDM BIT 15%.

### Base editing

Plasmids used in this study include pCMV_ABEmax_P2A_GFP (112101; Addgene, Watertown, MA) and pCMV-T7-ABEmax(7.10)-SpRY-P2A-EGFP (140003; Addgene). *In vitro* transcription of mRNA from these plasmids was performed as previously described^15^. We manually designed single guide RNAs (sgRNAs) to reproduce the p.Met41Thr c.122T>C mutation (supplemental Table 1). We used chemically modified synthetic sgRNAs harboring 2′-O-methyl analogs and 3′-phosphorothioate nonhydrolyzable linkages at the first three 5′ and 3′ nucleotides (Synthego, Redwood City, CA). 2 × 10^5^ to 1 x 10^6^ HSPCs per condition were electroporated two days after thawing with 3.0 μg of the ABE-encoding mRNA and 3.2 μg of the synthetic sgRNA using the P3 primary cell 4D-Nucleofector X Kit S (Lonza, Basel, Switzerland) and the CA-137 program (Nucleofector 4D). Editing efficiency was assessed by genomic DNA extraction (DNA micro kit, 56304; Qiagen), PCR amplification of the region interest (primers are given in supplemental table 1), Sanger sequencing and EditR analysis^16^. For TP53 invalidation by adenine base editing, sgRNAs disturbing splice sites were designed with the help of the SpliceR v.1.3.0 software^17^ and screened in HUDEP-2 by DNA sequencing and western blot assessment of TP53 downregulation. sgRNATP53 (supplementary table 1) was selected as the more efficient sgRNA. In rescue experiments, 3.2 µg of sgRNA3 and 3.2 µg of sgRNATP53 were added to the nucleofection mix to edit both genes simultaneously.

### Ubiquitin-traps capture

Ubiquitin traps (TUBEs) capture was performed as previously described^18,19^. Briefly, for each condition 5×10^6^ cells were used. Cells were pretreated with bortezomib 20 nM for 4 hours and washed two times in PBS before being collected and lysed in TUBE buffer (50 mM sodium fluoride, 5 mM tetrasodic pyrophosphate, 10 mM β-glyceropyrophosphate, 1% Igepal CA-630, 2 mM EDTA, 20 mM Na2HPO4, 20 mM NaH2PO4, 1 mM Pefablock, bortezomib 40 nM, bafilomycine 40 nM, 1.2 mg/ml protease inhibitor cocktail; Roche, Basel, Switzerland), in the presence of 100 ug of TUBEs (Bmolecular, Toulouse, France) or GST control. Lysates were centrifugated at 4°C during 10min at 13,000 rpm and supernatants recovered (input fraction). Lysates were incubated overnight at 4°C with 100 µL of glutathione agarose beads. The next day, 50 µL of flowthrough fractions were collected from the supernatant, and Glutathione beads were washed three times with 1mL of PBS. The bound fraction was eluted using boiling buffer (Laemmli 3X, beta-mercaptoethanol 5%) and analysed by western blotting.

### Immunoprecipitation

For each condition 5 million cells were used. Cells were pretreated with bortezomib 20 nM for 4 hours and washed two times in PBS before being collected and lysed in TUBE buffer supplemented with 100 µg of TUBEs to protect ubiquitylated proteins from deconjugation/degradation. Lysates were cleared by centrifugation at 13000rpm for 15min at 4°C and extracts of equal amount of total protein were used to immunoprecipitated p53 using 2 μg of antibody (DO-1, Santa Cruz) and 20 μL of protein A magnetic beads (prewashed with lysis buffer) during 2h at 4°C. Next day beads were washed 3 times with 500 μL of lysis buffer/wash and using a magnetic rack holder. Immuno-purified material was eluted in 100 μL of boiling buffer and boiled for 5min. Immunoprecipitates were analyzed by Western blot. before western-blot analysis.

### Western blot

Equal number of cells were pelleted and immediately lysed in boiling buffer and boiled 5 minutes at 95°C. Samples were cast in 12% SDS-polyacrylamide gels (Biorad) and separated by an hour of migration at 120 V before wet transfer on PVDF membranes overnight at 4°C and 40 mA. The next day, the membrane was incubated with 5% fat-free milk before a two-hour incubation with primary antibodies. After three 15 minutes washes in PBS Tween 0.1%, the membrane was incubated with secondary antibodies for one hour. After three additional washes, the membrane was revealed with the Supersignal West Atto ultimate sensitivity substrate (Thermo scientific, A38555) on a ChemiDoc MP Imaging System (BioRad, 12003154). The signal was quantified by the Image Lab Touch software (BioRad, 12014300). Antibodies used in this study include: anti-p53 (DOI-1, Santa Cruz), anti-ubiquitin (P4D1, Santa Cruz), anti-P21 (Cell signaling), anti-MDM2 (IF2, Thermofischer scientific).

### Isolation of mRNAs and RNA-sequencing

RNA was extracted with the Nucleospin RNA kit (Macherey-Nagel, 740955.50). Ribosomal RNA removal, RNA fragmentation, cDNA synthesis, end repair, 3’adenylation, adaptor ligation, PCR amplification, circularization, DNA nanoball synthesis, and sequencing on DNBSEQ (DNBSEQ Technology PE100) platform were performed by Beijing Genomics Institute (BGI, Hong Kong).

For RNA-seq data analysis, raw reads were subjected to quality control and adapter trimming using Trimgalore (version 0.6.6). Trimmed reads were then aligned to the reference genome (hg19) usingSTAR aligner (version 2.7.9a)^20^. Converting sam to bam, sorting and indexing bam files have been performed using samtools^21^ (version 1.11). Aligned sequencing reads have been count using HTSeq tool^22^ (version 0.11.3; options: -igene_id --additional-attr gene_name-t exon). Differentially expressed genes analysis have been detected using DESeq2 R package (version 1.38.3)^23^. Bigwig files have been generated using bamCoverage tool from deepTools suite^24^ (version 3.4.2;options: --smoothLength 15 --normalizeUsing RPKM). For gene set enrichment analysis, GSEA application (version 4.3.3) has been used^25^. Enrichment analysis has been performed using Enrichr R packages^26^.

### Isolation of total RNAs and northern blot analyses

For total RNA isolation, 300 000 to 10 × 10^6^ cells were collected by centrifugation for 5 min at 300 g and 4°C. After rinsing with 10 ml cold PBS, each pellet was thoroughly mixed with 1 ml TRI reagent and extracted following the manufacturer’s instructions. The upper aqueous phase was further purified by phenol/chloroform/isoamyl alcohol (25:24:1) extractions (Sigma) prior to isopropanol precipitation. Quantifications were performed at 260 nm with an Implen NP80 spectrophotometer (Mettler Toledo). Pre-rRNA precursors were separated on a 1.1% agarose gel containing 1.2% formaldehyde and 1× Tri/Tri buffer (30 mM triethanolamine, 30 mM tricine, pH 7.9) (3 µg total RNAs/lane). RNAs were transferred to Hybond N + nylon membrane (GE Healthcare) and cross-linked under UV light. Membrane pre-hybridization for 2 hours in 6× SSC, 5× Denhardt’s solution, 0.5% SDS, 0.9 µg/ml tRNA and then incubated overnight with the corresponding 5’-radiolabeled oligonucleotide probe. After washing twice for 10 min in 2×SSC, 0.1% SDS and once in 1× SSC, 0.1% SDS, the membrane was exposed to a PhosphorImager screen. Radioactive signals were revealed using a Typhoon Trio PhosphorImager (Cytivia) and quantified using the ImageQuantTL software. The probes used are listed in Supplementary Table 1.

### Transmission electron microscopy

Cells were first fixed 1.5h at room temperature with 2x fixation solution (5% glutaraldehyde and 4% paraformaldehyde (EMS, Delta-Microscopies, France) in 0.1 M Sorensen buffer, pH 7.2) for 15 min and then with 1x fixation solution (2.5% glutaraldehyde and 2% paraformaldehyde in 0.1 M Sorensen buffer, pH 7.2) overnight at 4°C. After 3 washes in 0.1 M Sorensen buffer, post-fixation was performed at RT with 1% OsO4 in 0.1 M cacodylate buffer, pH 7.2 buffer. Cells were pelleted, concentrated in Agar and treated for 1 h with 1% aqueous uranyl acetate. The samples were then dehydrated in a graded ethanol series and embedded in Epon (EMBed-812, EMS). After 48 h of polymerization at 60 °C, ultrathin sections (80 nm thick) were mounted on 200 mesh Formvar-carbon-coated copper grids. Finally, sections were stained with Uranyless and lead citrate (em-grade.com). Grids were examined with a TEM (Jeol JEM-1400, JEOL Ltd, Tokyo, Japan) at 80 kV. Images were acquired using a digital camera (Gatan Rio9, Gatan Inc, Pleasanton, CA, USA).

## Results

### Red blood cell Analysis in VEXAS Patients

In order to determine the impact of *UBA1* variants on red cells, we isolated mature RBCs from eight VEXAS patients with various degrees of anemia, and compared them with age and sex-matched controls (Table 1). We observed no significant differences between the two groups, either morphologically in mean projected surface area (PSA) as assessed by imaging flow cytometry (p = 0.40, figure 1A, n = 8), or in the prevalence of lactadherin-positive erythrocytes and reticulocytes (p = 0.76 and p = 0.77, figure 1B-C), or in deformability, evaluated using LORRCA (p = 0.36, figure 1D). Given the roles of both the abnormal presence of mitochondria in RBCs in several inflammatory conditions and of the ubiquitin-proteasome system in mitochondrial clearance during erythropoiesis^6^, we investigated mitochondrial retention, which may have, at least in part, explained inflammation. Mitotracker Deep Red staining indicated no differences in mitochondrial content between patients and controls (p = 0.95, figure 1E). Western blot analysis of total ubiquitin showed similar levels of polyubiquitylated proteins in both groups (p > 0.99, figure 1F, supplementary 1A). Proteomic analysis of RBCs from five patients (P1, P2, P5, P6, P7) and five controls demonstrated high correlation in proteome profiles (r² > 0.89 for each comparison, Figure 1G), except that VEXAS RBCs exhibited higher gamma-globin expression (p < 10⁻³, supplementary figure 1B). Thus, overall, RBC morphology, deformability, polyubiquitin content, and proteome did not show significant alterations between VEXAS patients and controls. These findings may suggest that mut-*UBA1* has no impact on RBC biogenesis and that inflammation is solely due to abnormal myeloid cells as reported earlier^27^. Alternatively, erythropoiesis may be defective and/or inefficient in mut-*UBA1* erythroid progenitors/precursors.

**Figure 1:**
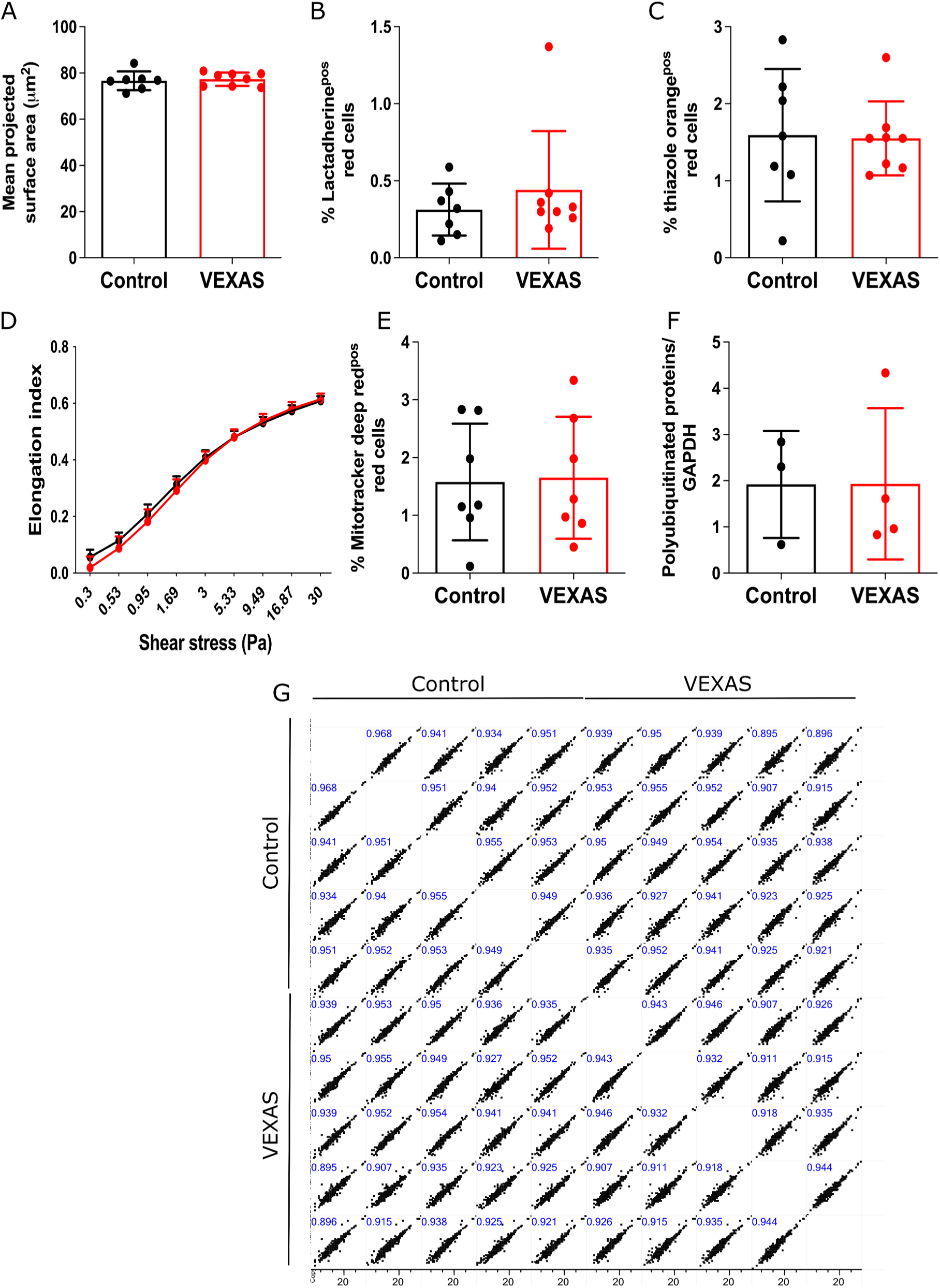
Characterization of red blood cells (RBCs) in VEXAS patients. A: histograms of the mean projected surface volume (µm^2^) of red cells isolated from patients and controls (n = 8). B: histograms of the percentage of lactadherine positive red cells determined by flow cytometry (n = 8). C: histograms of the percentage of thiazole orange positive red cells determined by flow cytometry (n = 7). D: a LORRCA rheometer was used to evaluate the RBC elongation index at various shear stresses in control (black) and VEXAS (red). Data are presented as mean +/- SD (n = 8). E: histograms of the percentage of red cells with mitochondrial retention, assessed by the positivity of Mitotracker Deep Red staining in flow cytometry (n = 8). F: quantification of the relative intensity of the polyubiquitin smear to the GAPDH band in western blot (n = 4). G: representation of proteomics results by a scatter plot of all RBC samples with r2 coefficients (n = 5).

**Table 1:**
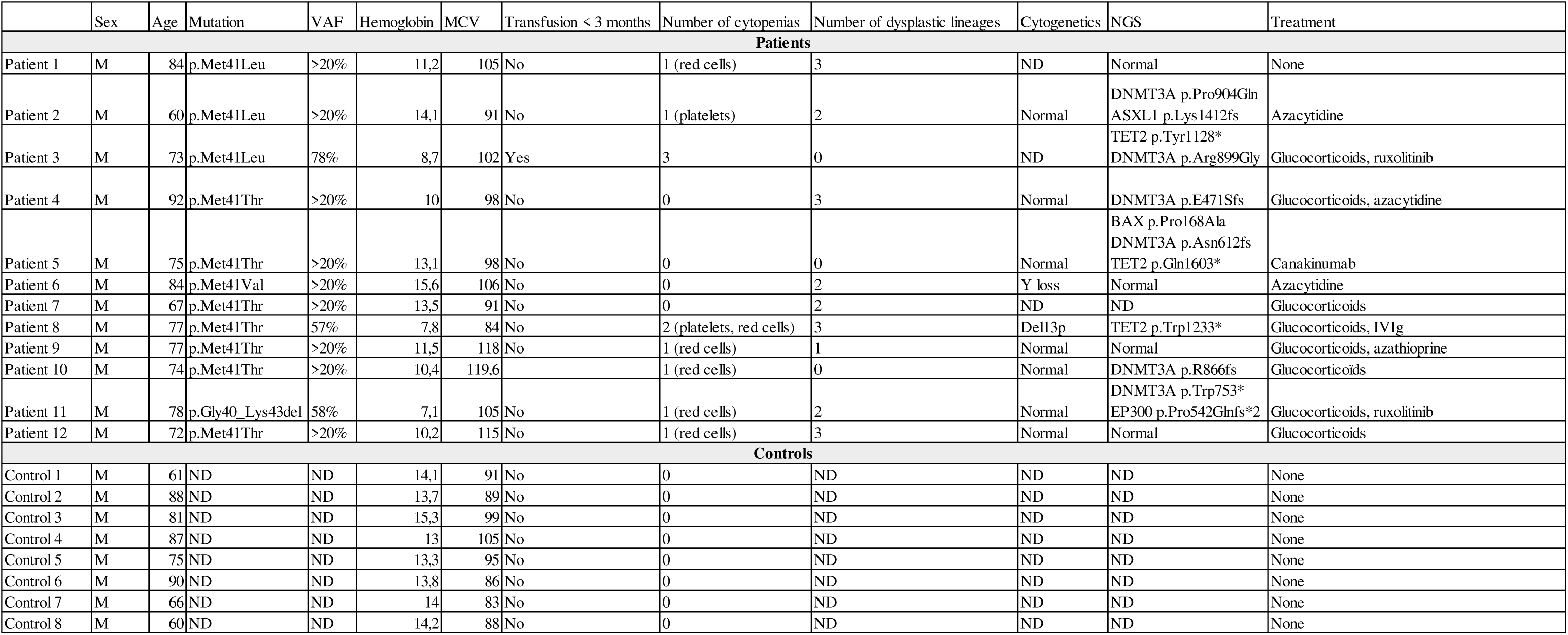
clinical and biological characteristics of patients and controls included in our study. Age, hemoglobin and MCV information are the parameters corresponding to the day patients were sampled for our study. *UBA1* VAF, when available, is the nearest measure to sampling day (+/- 3 months). Cytopenias are defined according to the following thresholds : hemoglobin < 13 g/dL; absolute neutrophil count < 1.8 G/L; platelets < 150 G/L. Cytogenetic anomality or clonal hematopoiesis detected at a VAF > 2% are reported if present at least once at any time during patient follow-up. Only VEXAS-targeting treatments or treatments affecting the erythroid lineage are reported. MCV : mean corpuscular volume. NGS : next generation sequencing. IVIg : intravenous immunoglobulin. ND : not done.

### Lack of Met41UBA1 Variant during in vivo and in vitro Erythropoiesis of VEXAS patients

In order to assess the contribution of mut-*UBA1* erythroid progenitors in erythropoiesis of VEXAS patients, we examined the allelic frequency of VEXAS-causing variants in the bone marrow of four patients. VEXAS variants were present in non-erythroid GPA^neg^ populations (VAF 31% ± 12.1, n = 4) but consistently absent in mature Band3^pos^CD49d^pos^ and Band3^pos^CD49d^neg^ populations (1% ± 2 and 1% ± 1, respectively, n = 3 and n = 4, Figures 2A-B). Then, we cultured peripheral blood CD34+ cells from four VEXAS patients with EPO and observed that VEXAS-causing variants present at day 0 disappeared by day 15 (Figure 2C-D). In two patients, the mutation persisted at day 7 in GPA^neg^ but not in GPA^pos^ populations, while it disappeared entirely in the others (Figure 2E). These findings suggest that VEXAS mutations may be lethal in the erythroid lineage, with erythropoiesis potentially rescued by the wt-*UBA1* hematopoietic population.

**Figure 2:**
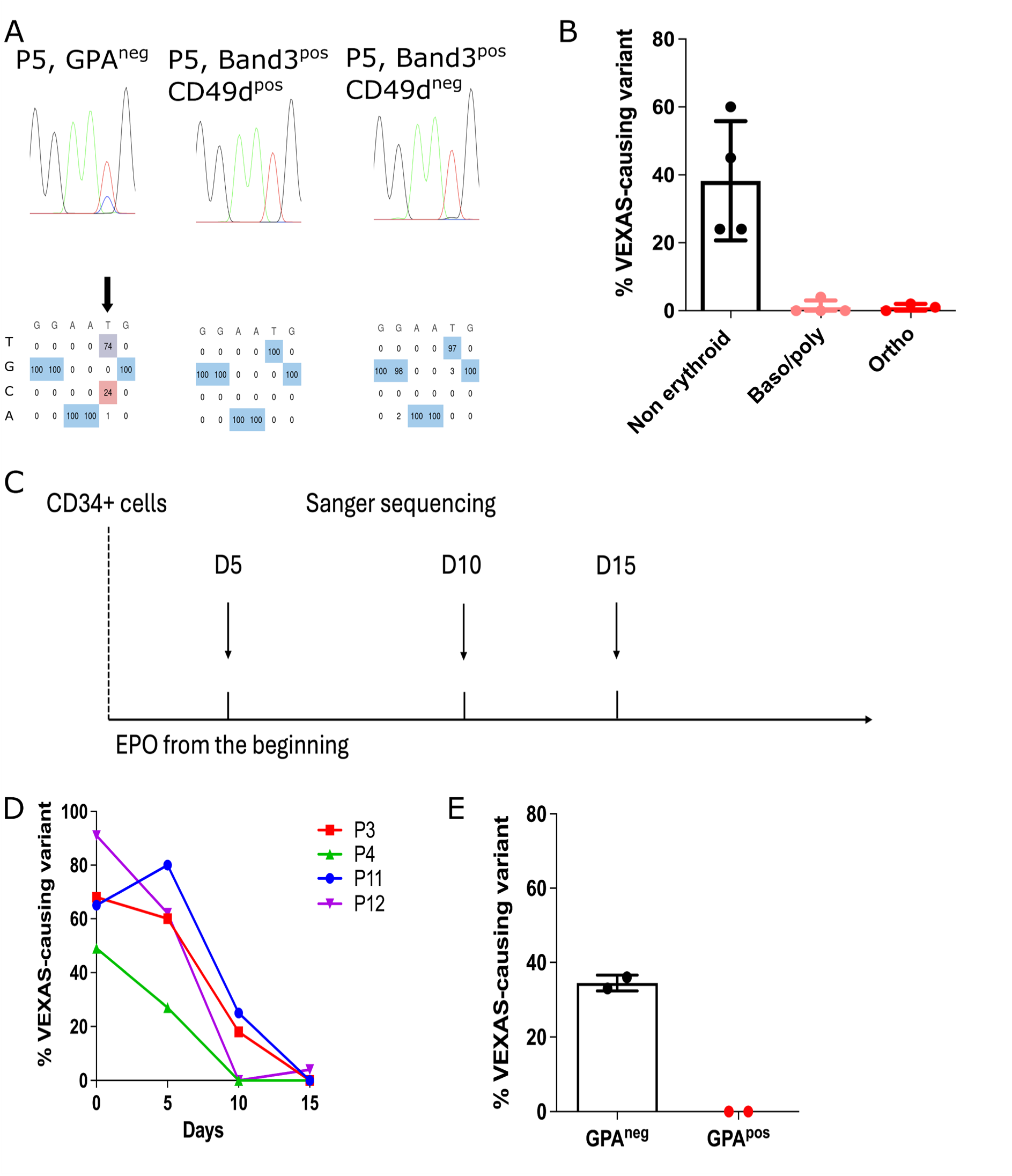
Absence of Met41 *UBA1* variants in VEXAS patients erythropoiesis. A: representative Sanger sequencing of *UBA1* exon 3 in different bone marrow populations in patient 5 (see table 1). B: graph showing the percentage of *UBA1*-mutated cells in non-erythroid bone marrow populations (GPA^neg^), basophilic and polychromatophilic erythroid cells (Band3^pos^CD49d^pos^), and orthochromatophilic erythroid cells (Band3^pos^CD49d^neg^), in four patients with VEXAS (P4, P5, P9, P10, see table 1). C: workflow of the erythroid culture of CD34+ isolated from the unmobilised peripheral blood (PB) of 4 VEXAS patients (P3, P5, P11, P12, see table 1). D: VAF of VEXAS-causing *UBA1* variants at day 0, 5, 10 and 15 of PB CD34+ erythroid cultures. E: histogram of the VAF of VEXAS-causing *UBA1* variants at day 7 of PB CD34+ culture in GPA^pos^ and GPA^neg^ populations in two patients (P3, P11).

### Impact of the Met41Thr UBA1 variant during in vitro erythropoiesis

To explore and explain the effects of VEXAS-causing variants, we introduced the p.Met41Thr c.122T>C variant into the HUDEP-2 erythroid cell line using adenine base editing. Four single guide RNAs (sgRNAs) effectively induced base editing, while sgRNA1 served as a control (Supplementary Figure 2A-B). The mutation disappeared after two weeks of maintenance culture with dexamethasone, with a mortality crisis during the second week post-electroporation (Supplementary Figure 2 C-D), particularly with sgRNA3, which had high base editing efficiency confirmed by a decrease in polyubiquitylated proteins (Supplementary Figure 2E) and vacuolization of apoptotic cells assessed by electron microscopy (Supplementary Figure 3). This suggests that the p.Met41Thr variant is lethal to HUDEP-2 cells at the pro-erythroblast stage. In human CD34^pos^ cord blood cells, introducing the p.Met41Thr c.122T>C variant with sgRNA3 resulted in proliferation arrest and massive cell death in CD36^pos^ cells (early erythroid progenitors) in the presence of SCF+Epo, but not in CD36^neg^ cells (early myeloid progenitors) in the presence of SCF+M-CSF (Figure 3A-F). In addition, erythroid differentiation was impaired by day 5 of phase II (post CD36 selection) in mut-*UBA1* cells, with virtually no GPA positive cells (Figure 3G). Moreover, the *UBA1* variant completely disappeared from erythroid live cells by day 12 (Figure 3H), but not in myeloid live cells in which the VAF decreased slowly. The latter observation is consistent with the presence of monocytopenia in patients^28^ and with reports of apoptosis in patients monocytic cultures^29^. When mut-*UBA1* cells were cultured with EPO without CD36 sorting, the proliferation defect and mortality crisis were less prominent (Supplementary Figure 4 A-B), further suggesting that the p.Met41Thr variant is less toxic in non-erythroid CD36^neg^ cells. A milder mortality crisis in non CD36-selected cells allowed us to sort GPA^pos^ cells for sequencing. The variant was present at a low VAF (mean 8% +/- 3.46, n = 3) in GPA^pos^ cells at day 7 of phase II, and absent from GPA^pos^ cells at day 12 (0 for the three replicates, supplementary Figure 4C). As a consequence, GPA expression of the bulk culture was diminished during the first week of culture, but not in the second week when wt-*UBA1* cells rescued erythropoiesis (Supplementary Figure 4D). These results align with observations in patients bone marrow, where only wt-*UBA1* erythropoiesis produces mature red cells and demonstrated that mut-*UBA1* is defective and not ineffective.

**Figure 3:**
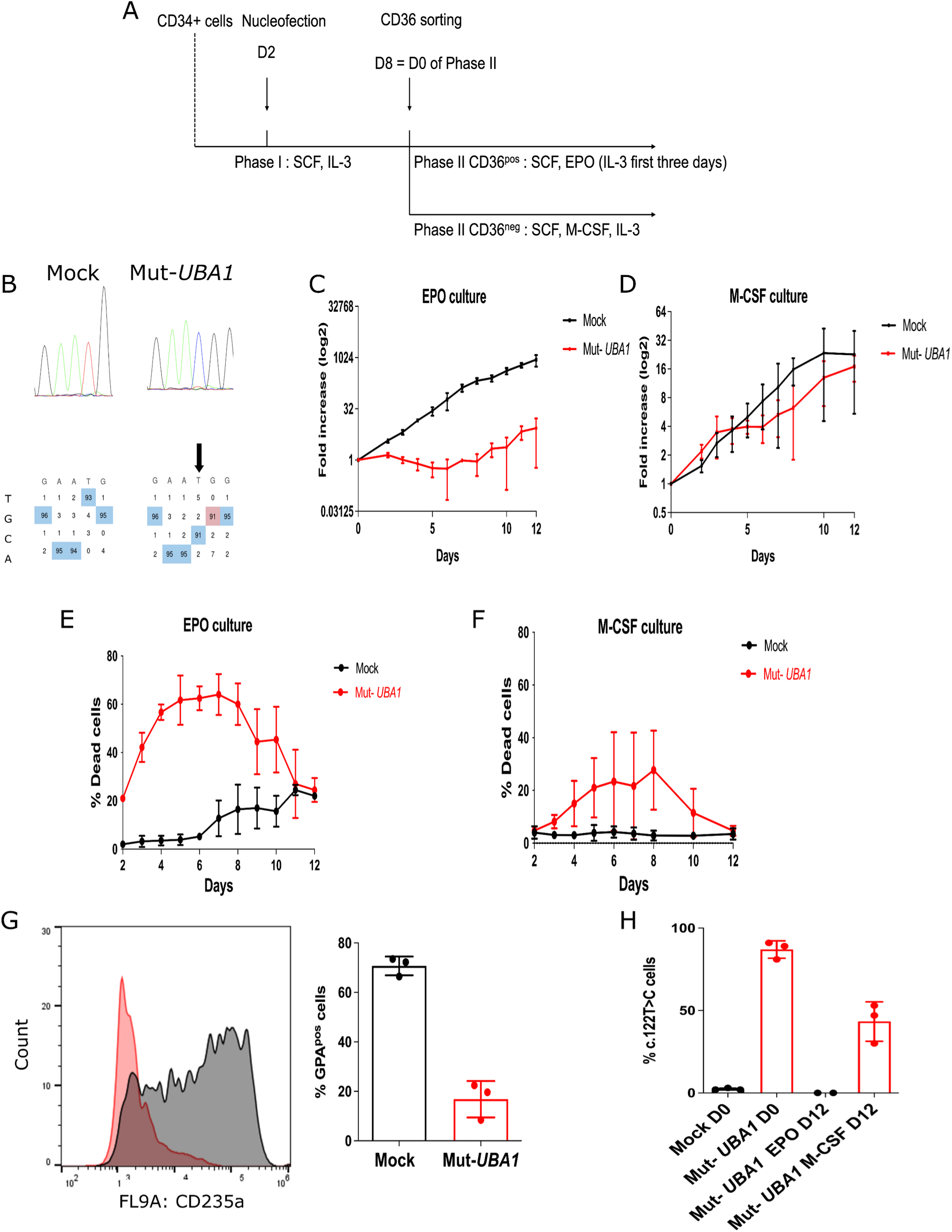
Impact of the p.Met41Thr variant on cord blood erythroid cultures. A: workflow of the editing procedure of cord blood CD34+ cells within our two-step erythroid culture. B: representative Sanger sequencing of *UBA1* exon 3 six days after electroporation (day 0 of phase II) of CD34+ cord blood cells with ABEmax alone (mock) or with ABEmax and SgRNA3 (mut-*UBA1*). C, D: fold-increase of mock and mut-*UBA1* cells during phase II, after CD36 sorting, in CD36^pos^ (C) and CD36^neg^ (D) cells (n = 3 different unrelated cord blood donors). E, F: daily mortality assessed by trypan blue staining in the aforementioned conditions. G: left: representative histogram of GPA expression in mock versus mut-*UBA1* cells at day 5 of phase II in the CD36^pos^ culture (n = 3 different unrelated cord blood donors). Right: histogram of the percentage of GPA^pos^ cells at day 5 of phase II in the CD36^pos^ culture (n = 3 different unrelated cord blood donors). H: histogram of p.Met41Thr VAF at day 0 (bulk sequencing) and at day 12 (sequencing of live propidium iodide negative cells) of phase II in the erythroid CD36^pos^ and myeloid CD36^neg^ cultures (n = 3 different unrelated cord blood donors).

### TP53 overexpression and abnormal ribosome biogenesis in mut-UBA1 erythroid cells

Then, mechanisms by which mut-*UBA1* erythropoiesis was defective were investigated. First, mut-*UBA1* cord blood cells died and could not proliferate as assessed by an increased number of 7-AAD and annexin V positive cells (supplementary Figure 5A). Because of the similitude between this observation and the early apoptosis of erythroid precursors due to haploinsufficiency in RPS19 or other ribosomal proteins leading to Diamond-Blackfan anemia (DBA), we investigated whether TP53 expression was increased and whether biogenesis of ribosomes was impaired. In agreement with these hypotheses, we observed a massive increase in TP53 expression in edited cells at day 0 of phase II of erythropoiesis (Figure 4A-B). Downstream transcriptional targets of TP53, namely P21 and HDM2 (human homolog of MDM2), were also overexpressed (Figure 4A), showing an activation of the TP53 pathway. RNA-seq comparing bulk mut-*UBA1* and mock cord blood cultures from three different biological donors at day 0 of phase II confirmed the transcriptional activation of the TP53 pathway in mut-*UBA1* cells (NES 2.40, p < 10^−4^, Figure 4C-D). Consistent with published transcriptomic data of VEXAS patients HSCs^27^, transcriptional activation of inflammatory and unfolded protein response (UPR) pathways was detected in mut-*UBA1* cord blood cells (supplementary figure 6 A-C), validating the relevance of our base-editing model. There was no difference in the levels of *TP53* mRNA between mut-*UBA1* and mock cord blood cells (p = 0.44, supplementary figure 5B), suggesting a post-transcriptional mechanism for TP53 overexpression. To understand this mechanism, we investigated the ubiquitylation of TP53. Both TUBE capture of ubiquitylated proteins revealed with an anti-TP53 antibody and immunoprecipitation of TP53 revealed with an anti-ubiquitin antibody showed that the increase of TP53 expression was associated with a reduction in its ubiquitylation (figure 4E-G).

**Figure 4:**
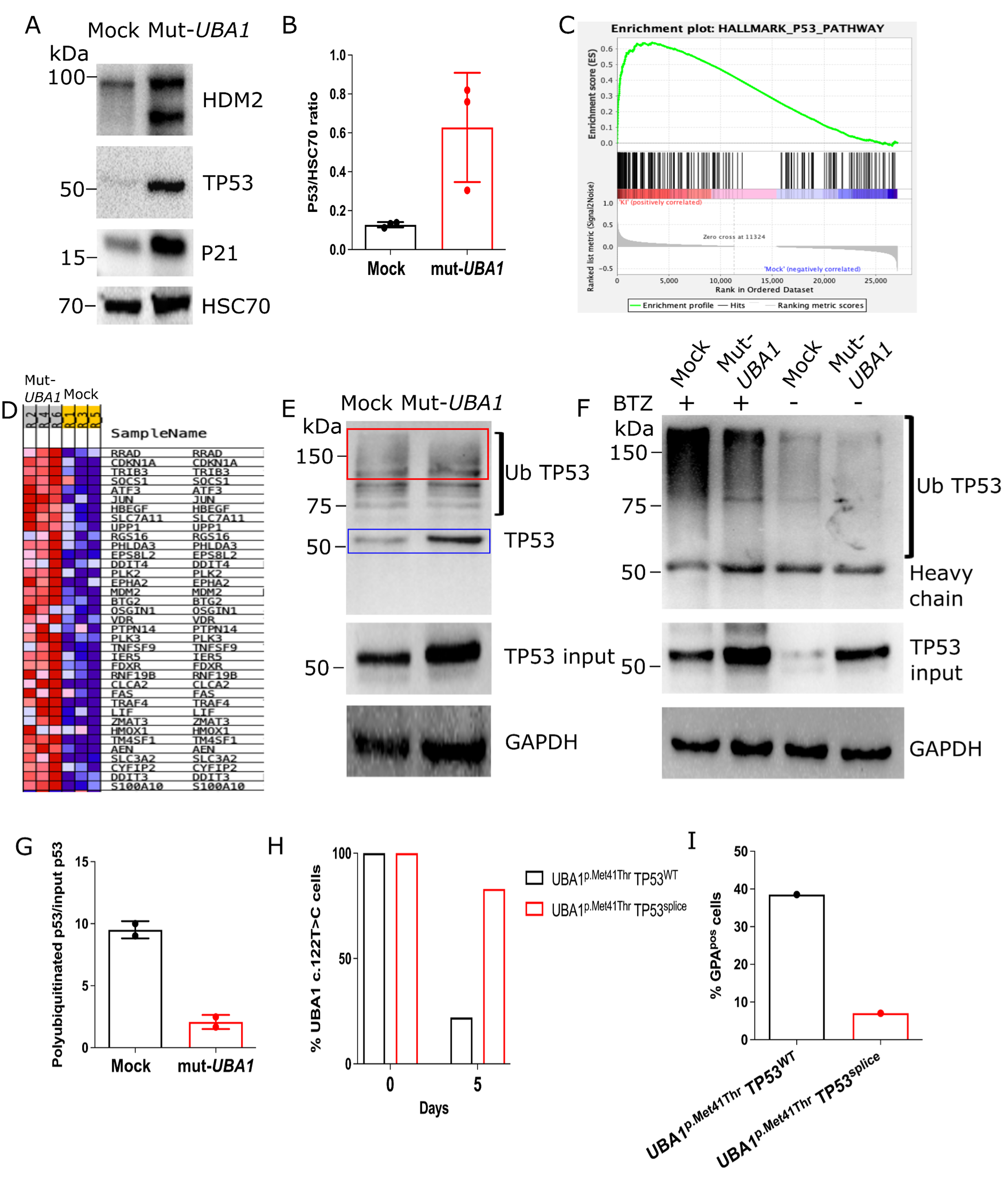
Defective ubiquitylation and overexpression of TP53 in mut-*UBA1* cord blood culture account for early erythroid lethality. A: representative western blot of TP53, P21 and HDM2 in mock or mut-*UBA1* cord blood cells at day 0 of phase II. B: quantification of TP53 expression compared to a loading control by western blot at day 0 of phase II in three different unrelated cord blood donors. C: GSEA analysis of the enrichment of transcripts from TP53 pathway genes in mut-*UBA1* cord blood cells at day 0 of phase II (n = 3 different unrelated donors). KI = mut-*UBA1*. D: Heat map of upregulated TP53 pathway transcripts in mut-*UBA1* cord blood cells (n = 3 different unrelated donors). E: representative experiment of a TUBE capture revealed against TP53 at day 0 of phase II in mock or mut-*UBA1* cord blood cells, with the loading control and TP53 revelation of the input. Red rectangle: fully polyubiquitylated TP53. Blue rectangle: non-ubiquitylated TP53 captured by TUBEs because one TP53 tetramer can contain both ubiquitylated and non-ubiquitylated units. F: representative immunoprecipitation of TP53 revealed with an anti-ubiquitin antibody, with the loading control (GAPDH) and TP53 revelation of the input, in mock or mut-*UBA1* cells treated or not with bortezomib (BTZ). (n = 2 different unrelated biological donors, both different from the donor displayed in D). G: quantification of the ubiquitylated TP53/input TP53 ratio in immunoprecipitation with BTZ pretreatment from two different unrelated cord blood donors. H: histogram of *UBA1* p.Met41Thr VAF in bulk culture (day 0) and in sorted live cells (annexin V-negative, propidium iodide negative, day 5 of phase II) in the CD36^pos^ culture, compared between cells solely edited for *UBA1* (UBA1^p.Met41Thr^ TP53^WT^) and cells edited for both *UBA1* and *TP53* (UBA1^p.Met41Thr^ TP53^splice^). I: histogram of the percentage of GPA^pos^ cells at day 5 of phase II in UBA1^p.Met41Thr^ P53^WT^ and UBA1^p.Met41Thr^ TP53^splice^ CD36^pos^ cells.

Since CD36^pos^ cells accounted for most of the increase in TP53 levels (supplementary Figure 5C), we hypothesized that, like as in DBA, TP53 could be a good candidate to explain erythroid-specific lethality of *UBA1* variants. Invalidation of TP53 expression by inducing the c.672+2T>C splice mutation in *TP53* (supplementary Figure 5D) with base editing increased survival of CD36^pos^ mut-*UBA1* cells (Figure 4H). Erythroid differentiation (Figure 4I) was however not rescued, suggesting that TP53 induction is not the only cause of erythropoiesis defect.

Then, we investigated if additional mechanisms could account for defective mut-*UBA1* erythropoiesis, and assessed ribosome biogenesis in mut-*UBA1* HUDEP-2 cells. In agreement with an alteration of ribosome biogenesis, the northern blot analysis of ribosomal RNAs showed an early cleavage defect at site 2, with accumulation of 47S, 41S and 43S precursors, decrease of downstream 30S, 32.5S and 21S precursors, and with the activation of an alternative maturation pathway with an increase in 36S levels (Figure 5A-B). However, the activation of the latter pathway was also defective, as 18S-E levels were not different between the two conditions. These findings suggest that the ribosome biogenesis defect in our model is global and affects several maturation events. Alteration of ribosome biogenesis was also supported by changes in the ultrastructure of the nucleoli in mut-*UBA1* cells: the dense fibrillar component (DFC) appeared thicker and more contrasted than in controls, while fibrillar centers (FC) were smaller or even difficult to distinguish (figure 5C).

**Figure 5:**
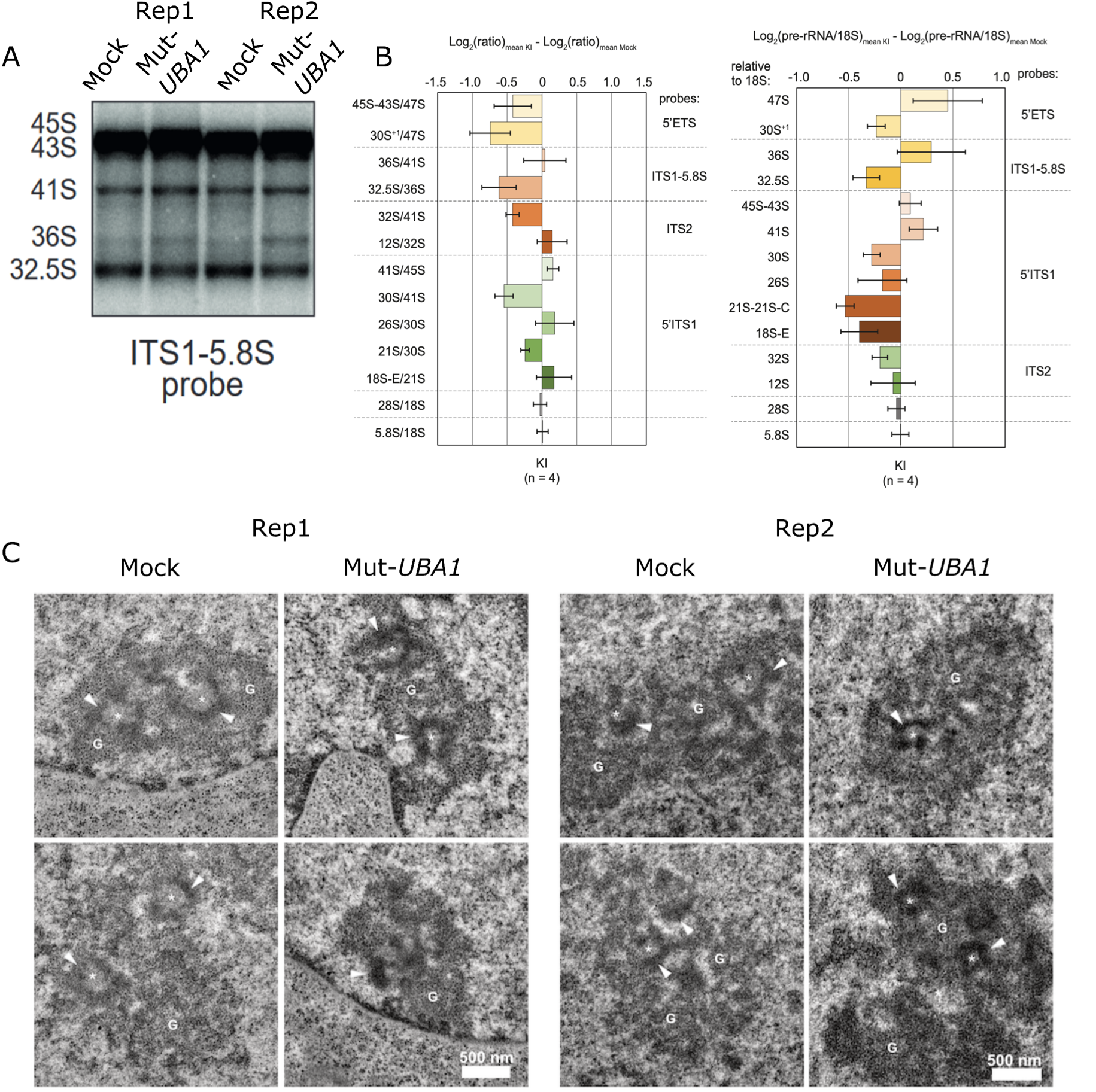
Alteration of ribosome biogenesis in mut-*UBA1* HUDEP-2 cells. A: representative northern blot revealed with the ITS1-5.8S probe showing increased level of 43S, 41S and 36S and decreased levels of 32.5S in mut-*UBA1* HUDEP-2 cells versus mock cells (n = 4 replicates). B: log2 ratios of product/precursor (left) or pre-rRNA/mature 18S rRNA (right) between mut-*UBA1* and mock HUDEP-2 cells (n = 4 replicates). C: changes in the ultrastructure of the nucleoli in mut-*UBA1* HUDEP-2 cells compared to control cells (n = 2 replicates). Grey levels were normalized in order to have comparable contrasts. Stars: fibrillar centers; Arrowheads: dense fibrillar component; G: granular component.

Overall, these findings show strong analogies between VEXAS and DBA and provide an explanation for the inability of mut-*UBA1* cells to complete erythroid differentiation.

### Targeting cellular vulnerabilities by bortezomib exposure of mut-UBA1 hematopoietic cells

Current treatments for VEXAS primarily aim to suppress inflammation by targeting myeloid cells and/or their inflammatory mediators, such as IL-6, or by interfering with downstream signaling pathways like JAK/STAT. Additionally, demethylating agents are used with the goal of improving erythropoiesis and restoring inhibitory inflammatory pathways in myeloid cells. Our findings suggest that, due to the mosaicism present in VEXAS (unlike in DBA), targeting cellular vulnerabilities may provide new therapeutic strategies to eliminate mut-*UBA1* cells and restore wt-*UBA1* hematopoiesis. As a proof-of-concept experiment, we used low doses of bortezomib, a proteasome inhibitor, to selectively induce apoptosis in mut-*UBA1* clones whose proteasome activity is already altered due to defective ubiquitylation, while preserving wt-*UBA1* hematopoiesis. Consistent with our hypothesis, 48 hours of treatment with bortezomib 10nM starting at day 2 of phase II led to an important cell death in CD36^neg^ mut-UBA1 cells cultured in the presence of M-CSF, but not in the CD36^neg^ mock cultures (Figures 6 A-C), underscoring its potential as a treatment option.

**Figure 6:**
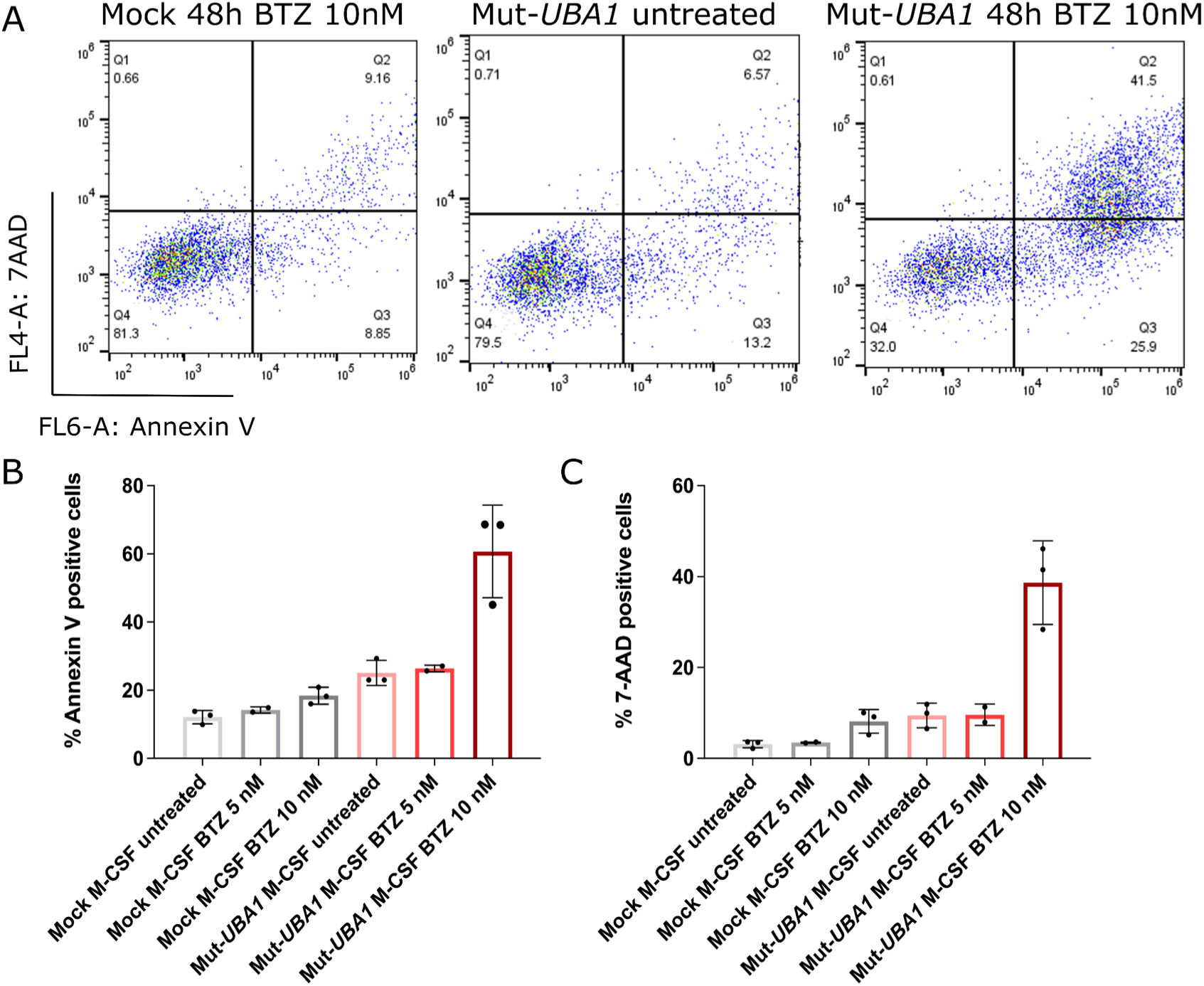
The p.Met41Thr variant sensitizes myeloid cord blood cells to cell death by bortezomib (BTZ), a proteasome inhibitor. A: representative flow cytometry analysis of annexin V/7-AAD staining 48 hours after continuous treatment with BTZ 10 nM in mut-*UBA1* cord blood CD36^neg^ cells, compared to staining of untreated mock and mut-*UBA1* CD36^neg^ cells. B: histograms of annexin V positivity without bortezomib or after 48 hours of 5nM or 10 nM bortezomib treatment (n = 3 different unrelated cord blood donors). C: histograms of 7-AAD positivity without bortezomib or after 48 hours of 5nM or 10 nM bortezomib treatment (n = 3 different unrelated cord blood donors).

## Discussion

In this study, we provide the first comprehensive characterization of red blood cells and erythroblasts in VEXAS syndrome. Our findings suggest that VEXAS represents a novel cause of mosaic erythroblastopenia, resembling Diamond-Blackfan Anemia (DBA), and that RBC production results from wt-*UBA1* erythropoiesis.

Firstly, utilizing state-of-the-art technologies, we have demonstrated that circulating RBCs in VEXAS patients neither exhibit significant abnormalities in shape or deformability compared to age-matched controls, nor do they contain residual mitochondria, which could have increased the risk of thrombosis and inflammation — a hallmark of the disease. The absence of difference in mean projected surface area between the patients and controls is, at first glance, surprising in the context of a macrocytic anemia, but to our knowledge no correlation between PSA and mean corpuscular volume (MCV) has been previously reported. In addition, to reduce the risk of confounding findings with physiological ageing, we chose to compare VEXAS patients with age-matched controls, in which prevalence of clonal hematopoiesis is high, and elevated MCV is a common sign of CH^30^.

Then, we found that protein ubiquitylation and protein content in patients RBCs were comparable to those of controls except for an increase in gamma-globin expression in VEXAS patients. Since elevated gamma-globin levels have been reported in various TP53-associated anemias (DBA, 5q del syndrome, Fanconi anemia)^31,32^ as well as in myelodysplastic syndromes and clonal hematopoiesis^33^, this finding does not necessarily reflect an ubiquitylation defect. It rather suggests that the mut-*UBA1* erythroid defect induces a wt-*UBA1* stress erythropoiesis or that clonal hematopoiesis and myelodysplastic clones are selected within the inflammatory bone marrow environment of VEXAS patients.

Secondly, sequencing of bone marrow erythroblasts from patients showed that *UBA1* mutations disappeared by the late basophilic/polychromatophilic stage, consistent with *in vitro* culture of peripheral blood CD34^pos^ hematopoietic stem cells from patients that showed extinction of *UBA1* variants during early erythropoiesis.

Thirdly, introducing the p.Met41Thr variant resulted in early lethality in pro-erythroblastic HUDEP-2 cell line and cord blood CD36^pos^ erythroid precursors, more important that the lethality observed in cord blood monocytic CD36^neg^ precursors cultured with M-CSF.

We thus propose that mut-*UBA1* erythroblasts undergo early apoptosis, with red cell production being compensated by wt-*UBA1* erythroblasts. This aligns with reports of erythroid hypoplasia in the bone marrow of VEXAS patients, as would be expected in a mosaic erythroblastopenia^34^. In our model the degree of anemia in VEXAS would depend on the ability of wt-*UBA1* bone marrow compartment to restore erythropoiesis. This capacity may depend on intrinsic parameters such as the selection of clonal hematopoiesis or myelodysplastic clones and/or extrinsic parameters such as an unfavorable inflammatory bone marrow environment or treatment with ruxolitinib. This model, however, does not fully explain why VEXAS patients without other hematological conditions exhibit macrocytosis. The latter might be due to clonal hematopoiesis in wt-*UBA1* compartment, early arrest in erythroblast divisions, or by induction of a stress erythropoiesis in the context of a TP53-associated anemia with insufficient compensation by wt-*UBA1* erythroblasts to restore normal hemoglobin levels. Indeed, macrocytosis is a feature of Diamond-Blackfan and Fanconi anemias^35,36^, as well as 5q del syndrome^37^. In the latter, 5qdel erythroblasts display a maturation arrest^37^, and as in VEXAS, most red cells originate from wild-type erythroblasts, yet displaying macrocytosis.

Our data also demonstrates that *in vitro* VEXAS erythropoiesis shares features with Diamond-Blackfan and Fanconi anemias, such as TP53 overexpression. Unlike Cas9 nucleases, adenine base editors do not activate the p53 pathway in human CD34^pos^ cells^38,39^, indicating that TP53 overexpression in mut-*UBA1* cells is specific to the mutation, and not a side-effect of base editing.

Interestingly, our data suggests that the cytoplasm-restricted ubiquitylation defect in VEXAS leads to reduced TP53 ubiquitylation, despite existing models suggesting that HDM2 can ubiquitylate TP53 in both the nucleus and cytoplasm^40,41^. We can formulate two hypotheses about this finding. First, excess ribosomal proteins, accumulated in the context of disturbed biogenesis or defective ubiquitylation and degradation, could bind HDM2 and inhibit its ubiquitin ligase activity in the nucleus where UBA1a function is conserved. Alternatively, cytoplasmic ubiquitylation could be a necessary condition for TP53 degradation, potentially to counteract deubiquitylases present in the cytoplasm such as HAUSP/USP7^42^. This would align with Shi et al.’s work identifying CBP as a cytoplasm-restricted ubiquitin ligase for TP53, where its knockdown reduces TP53 ubiquitylation^43^.

In previous reports it has been shown that erythroblasts are particularly sensitive to TP53-mediated cell death as well as proliferation and differentiation arrest^44^, which may explain the observed cell death in base-edited erythroid cultures. However, in our VEXAS model, the pilot experiment that invalidated TP53 expression could restore survival but not differentiation. This suggests TP53-independant mechanisms in VEXAS erythroblastopenia, at least for differentiation, similar to what has been described in some DBA genotypes^45,46^.

Bringing the analogy between VEXAS anemia and DBA to the next level, we showed that the p.Met41Thr variant induces defects in the early stages of ribosome biogenesis, even though this process takes place in the nucleus. This could be due to defective clearance of abnormal mature ribosomes and subsequent impaired recycling of ribosomal proteins from the cytoplasm to the nucleus, especially in the event of collided ribosomes^47^. The defects in ribosomal biogenesis further explain the toxicity of VEXAS variants in erythroblasts, but also unravel that a cytoplasmic ubiquitylation defect can have consequences in the nuclear compartment. Our base-editing model of VEXAS syndrome is thus a new tool to investigate the mechanisms of ubiquitylation of shuttling, nucleocytoplasmic proteins as well as the impact of cytoplasmic ubiquitylation on nuclear events.

Finally, in a proof-of-concept experiment, we show that our model can be used to screen drugs aiming at killing myeloid mut-*UBA1* cells without affecting wild-type cells. As mutated cells are already weakened by proteotoxic and ribosomal stress, bortezomib, a proteasome inhibitor usually toxic in the lymphoid lineage, was able to induce specific apoptosis of myeloid mutated cells *in vitro* at low dose without affecting the wild-type control cells, which may reduce inflammation, allowing a full potential of wt-*UBA-1* erythropoiesis. The fact that VEXAS is not associated with leukemic transformation should encourage researchers and clinicians to find mild cytotoxic agents that exploit the molecular vulnerability of *UBA1* clones. While new treatments should be developed to eliminate the mut-*UBA1* myeloid clones responsible for inflammation, we propose that the treatment of anemia in VEXAS should instead focus on the wt-*UBA1* compartment. First of all, severe anemia in the context of VEXAS should prompt investigation for additional hematopoietic conditions such as a myelodysplastic syndrome to account for insufficient compensation of the mosaic erythroblastopenia by the wt-*UBA1* erythroid compartment. Then, tailored treatments should aim at restoring wt-*UBA1* erythropoiesis with erythroid stimulating agents like erythropoietin, or, in case of an associated myelodysplastic syndrome, with lenalidomide, 5-azacytidine or TGF-beta ligand traps, depending on the additional molecular abnormalities in the wt-*UBA1* clones.

In conclusion, we identify VEXAS anemia as a mosaic erythroblastopenia, highlighting the crucial role of cytoplasmic ubiquitylation in regulating TP53 protein levels as well as ribosome biogenesis. This discovery also provides meaningful insights for clinicians managing VEXAS patients.

## Authors contributions

FR conceived the study, designed and performed most experiments, recruited most patients, interpreted results and wrote the manuscript. GH and A Miccio designed base editing experiments. SEH designed erythropoiesis experiments and interpreted results. AG and MD helped in sequencing experiments. EFG designed, performed and interpreted proteomics experiments. MD, SP and PA designed, performed and interpreted AMNIS and LORRCA experiments. VS and PEG designed, performed and interpreted transmission electron microscopy experiments. TT and JR secured funding. VJ, A Mekinian, SGL, and OK recruited patients and provided clinical information. EA helped in RNA isolation. MS and ES designed the RNA-seq experiments, analyzed the raw data and interpreted RNA-seq results. MFO designed, performed and interpreted northern blot experiments. MR designed, performed and interpretated TP53 ubiquitylation experiments. OH conceived and designed the study, interpreted results, wrote the manuscript and secured funding. All authors revised the manuscript and approved its final version.

## Acknowledgments

We thank Drs. Eric Bouvard, Katayoun Jondeau, Jean-Benoît Arlet, Edouard Flamarion, Mathilde Devaux, Thomas Hanslik for patient recruitment. We thank Dr. Erika Brunet for help in pilot unpublished homology directed repair (HDR) experiments. We thank the CRB (centre de ressources biologiques) of hôpital Saint-Louis for processing and providing cord blood samples. We thank Dr. Jérôme Megret from the SFR Necker Cytometry platform for help in cell sorting. We acknowledge the METi imaging facility, member of Genotoul and of the national infrastructure France-BioImaging supported by the French National Research Agency (ANR-10-INBS-04). FR was supported by a PhD scholarship from the Laboratory of Excellence GR-Ex. MD holds a Paris-Cité University research engineer position fully supported by the Laboratory of Excellence GR-Ex.

## Disclosures

**Thiago Trovati Maciel:** *LGD France:* Research Funding; *Imara Inc.:* Research Funding; *Alexion Pharmaceuticals:* Research Funding; *Bristol-Myers Squibb:* Research Funding; *F. Hoffmann-La Roche Ltd:* Research Funding. **Manuel Rodriguez:** *Bmolecular:* Consultancy, Current equity holder in publicly-traded company. **Olivier Hermine:** *AB Science:* Consultancy, Current equity holder in publicly-traded company, Patents & Royalties, Research Funding; *Inatherys:* Consultancy, Current equity holder in publicly-traded company, Patents & Royalties, Research Funding; *BMS:* Research Funding; *Alexion:* Research Funding; *Roche:* Research Funding; *MSD Avenir:* Research Funding.

## Notes

### Competing Interest Statement

This study was funded by MSD.

