## Supplementary material for "VEXAS anemia is a mosaic erythroblastopenia"

**Supplementary figure 1**

A B

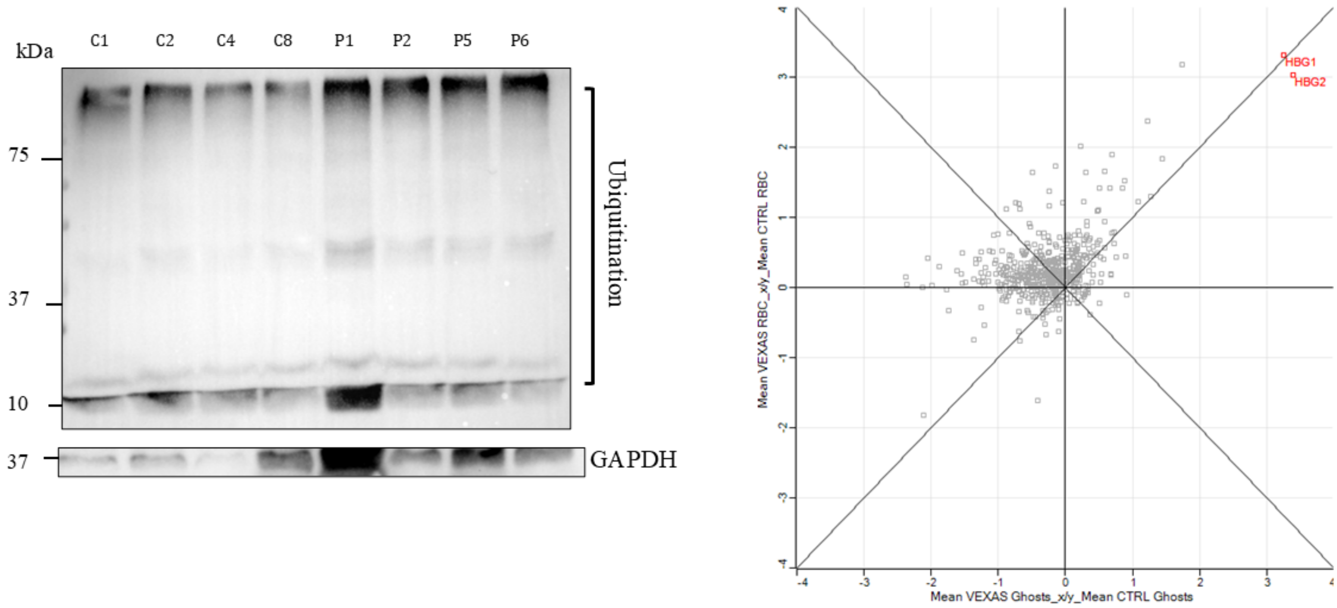


**Supplementary Figure 1:** Further characterization of VEXAS RBCs. A: representative Western blot of ubiquitin and GAPDH in RBCs of patients and controls, with 5 million sorted mature RBCs loaded per lane. The upper smear is the signal from polyubiquitinated proteins, the lower smear from free ubiquitin. B: representation of proteins quantified in red cells in at least 3 VEXAS patients and 3 controls as a VEXAS/control log2 ratio in total red cells (left axis) or in ghosts (bottom axis). Proteins statistically differentially expressed between VEXAS patients and controls (p-value < 0.01, log2 ratio > 1) are labeled in red.

#### Supplementary figure 2

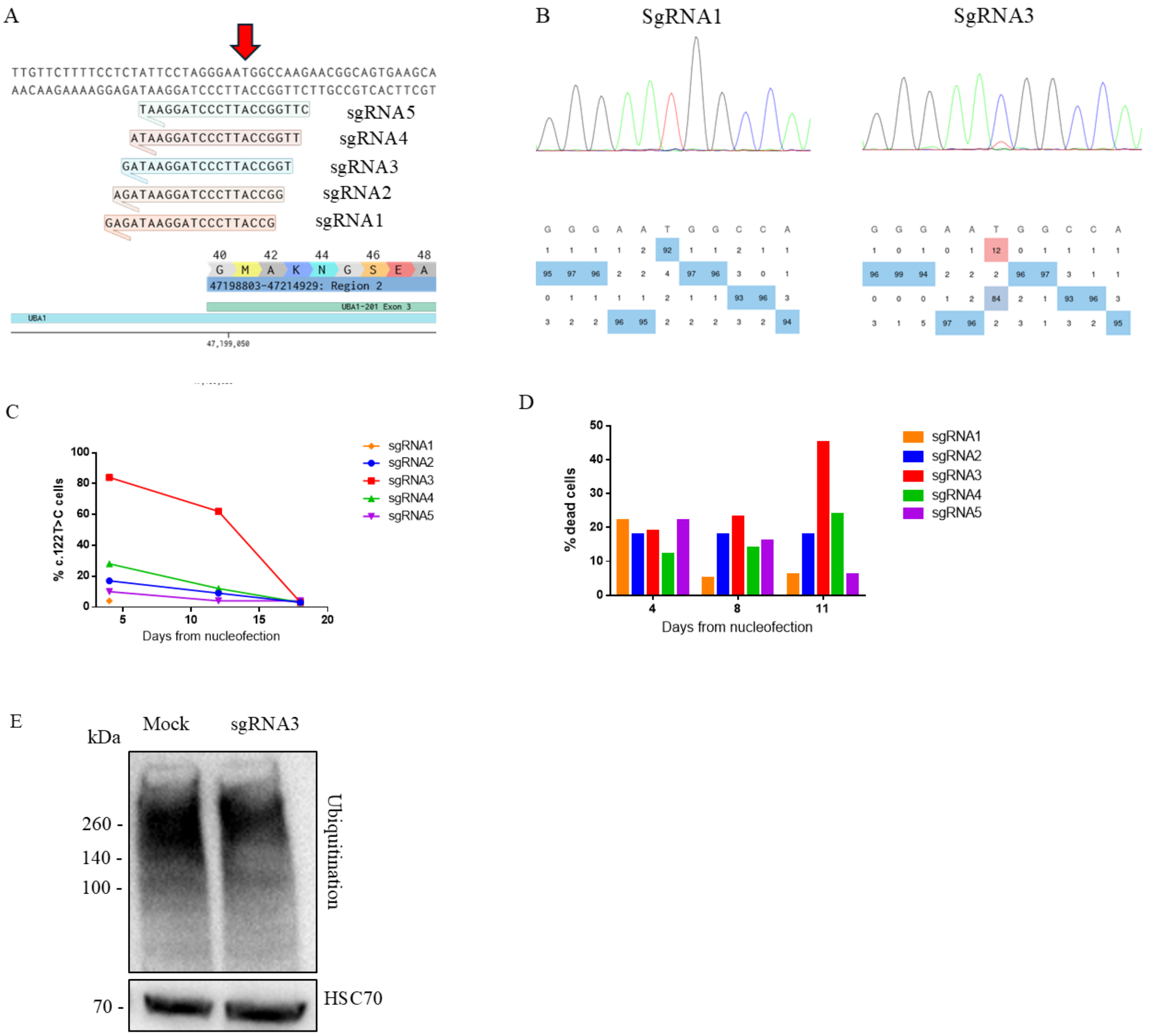


**Supplementary figure 2**: **Impact of the p.Met41Thr variant on a maintenance culture of HUDEP-2 cells.** A: design of 5 sgRNAs for adenine base editing of the p.Met41Thr mutation in HUDEP-2 cells. B: representative Sanger sequencing of the exon 3 of *UBA1* 4 days after electroporation with sgRNA1 (control) or sgRNA3. C: VAF of *UBA1* c.122T>C at days 4, 11 and 18 of an amplification culture of HUDEP-2 edited with sgRNAs 1-5. D: trypan blue-assessed mortality of HUDEP-2 cells at day 4, 11 and 18 of nucleofection with sgRNAs 1-5. E: western blot of polyubiquitinated proteins six days after nucleofection in HUDEP-2 cells edited with sgRNA3 compared to mock HUDEP-2 cells, electroporated with ABEmax without sgRNA.

**Supplementary figure 3**

A Mock

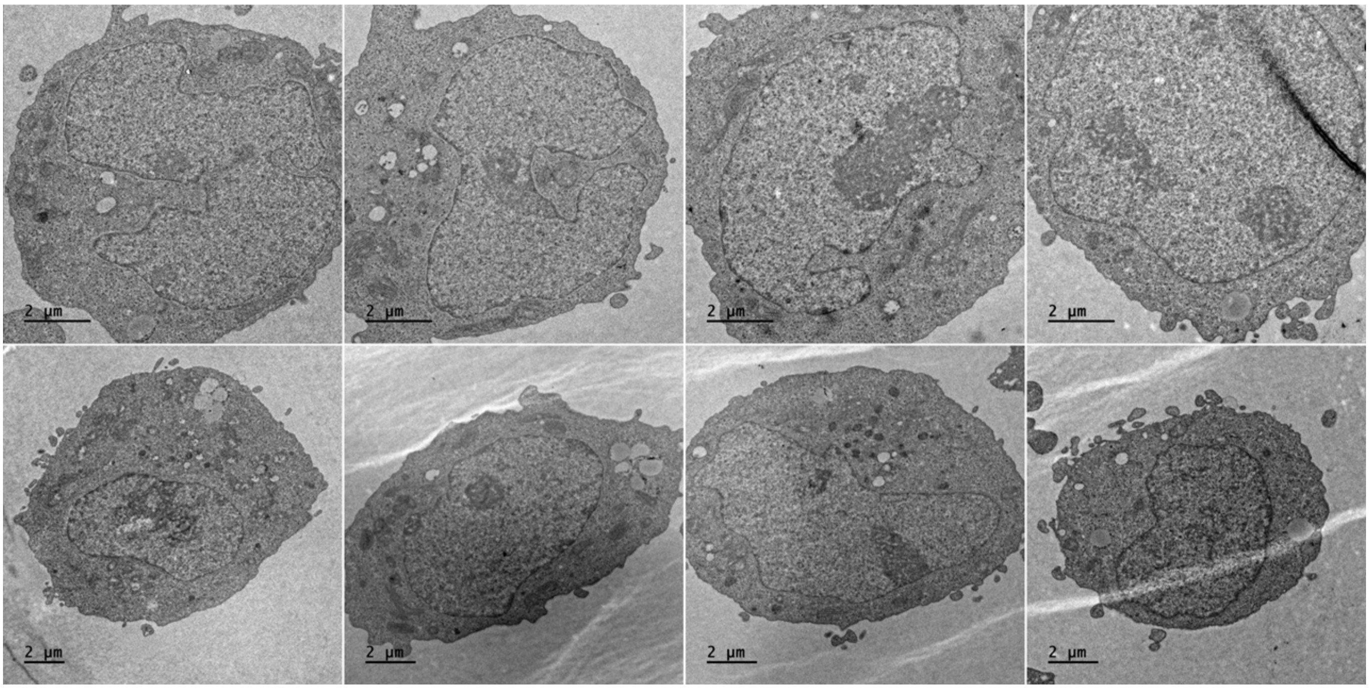


B Mut-*UBA1*

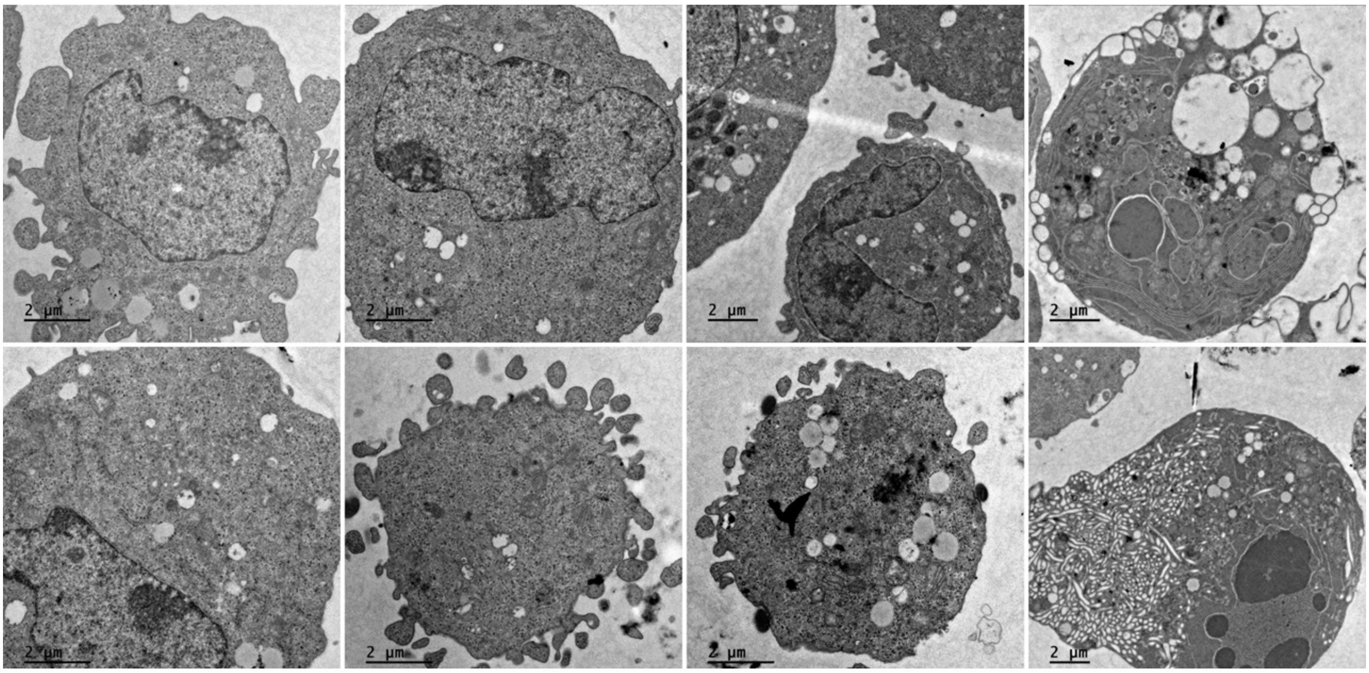


**Supplementary figure 3: Ultrastructure of mut-*UBA1* HUDEP-2 cells.** A: Transmission electron microscopy (TEM) broad views of mock HUDEP-2 cells six days after electroporation with ABEmax but without sgRNA. B: TEM broad views of mut-*UBA1* HUDEP-2 cells six days after electroporation with ABEmax and sgRNA3.

**Supplementary figure 4**

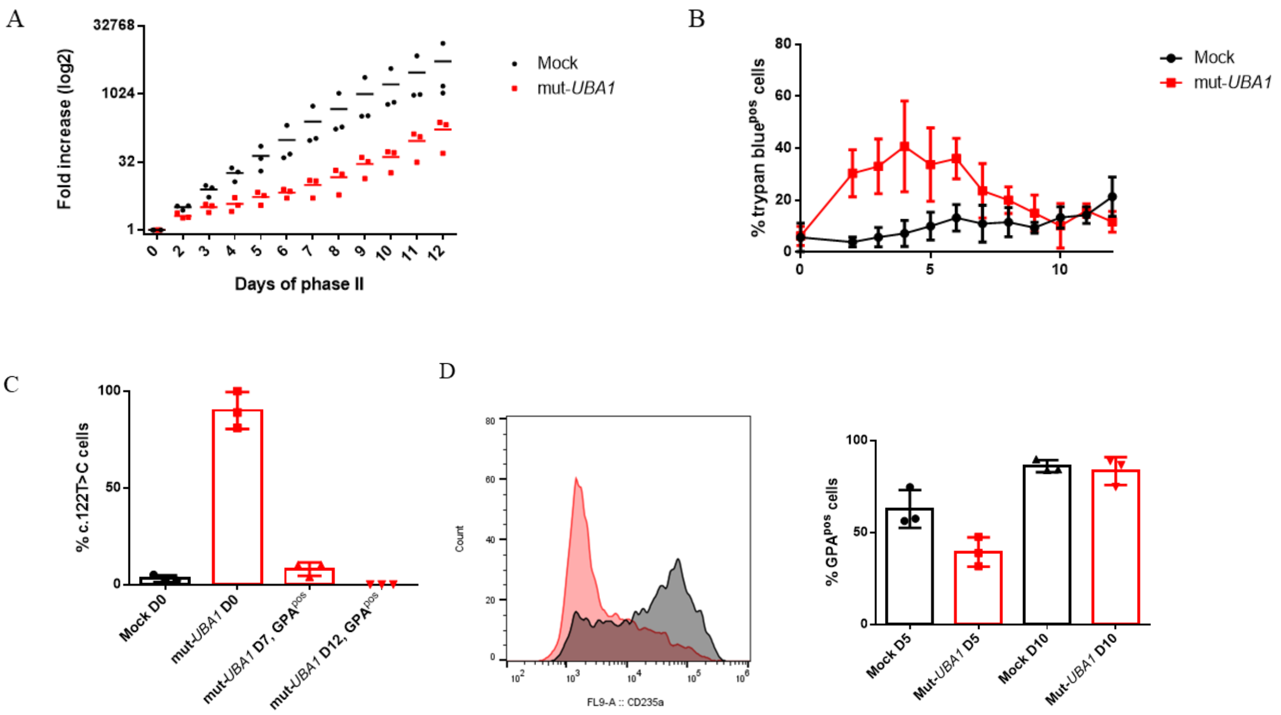


**Supplementary figure 4**: **Impact of the p.Met41Thr variant on cord blood erythroid cultures without CD36 sorting.** A: fold-increase of mut-*UBA1* vs mock cord blood cells cultured with EPO without CD36 sorting during phase II (n = 3 different unrelated donors). B: daily mortality assessed by trypan blue staining in mut-*UBA1* vs mock cord blood cells cultured with EPO without CD36 sorting during phase II (n = 3 different unrelated donors). C: histograms of c.122T>C VAF in the bulk culture (day 0) or in GPA$^{pos}$ sorted cells (days 7 and 12) in the EPO culture without CD36 sorting (n = 3 different unrelated cord blood donors). D: left: representative histogram of GPA expression at day 5 of phase II in mock (black) or mut-UBA1 (red) cord blood cells without CD36 sorting. Right: histograms of the percentage of GPA$^{pos}$ cells at days 5 and 10 of phase II in mut-*UBA1* vs mock cord blood cells cultured with EPO and without CD36 sorting (n = 3 different unrelated donors).

### Supplementary figure 5

A

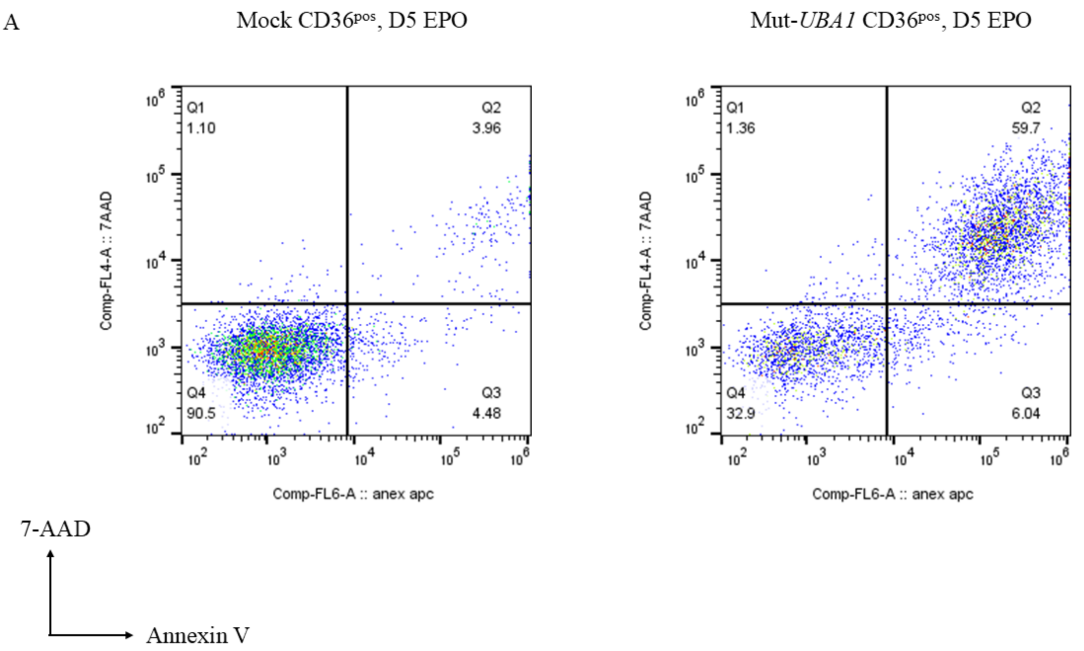


B

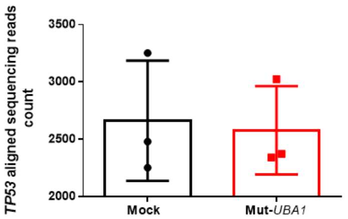


C

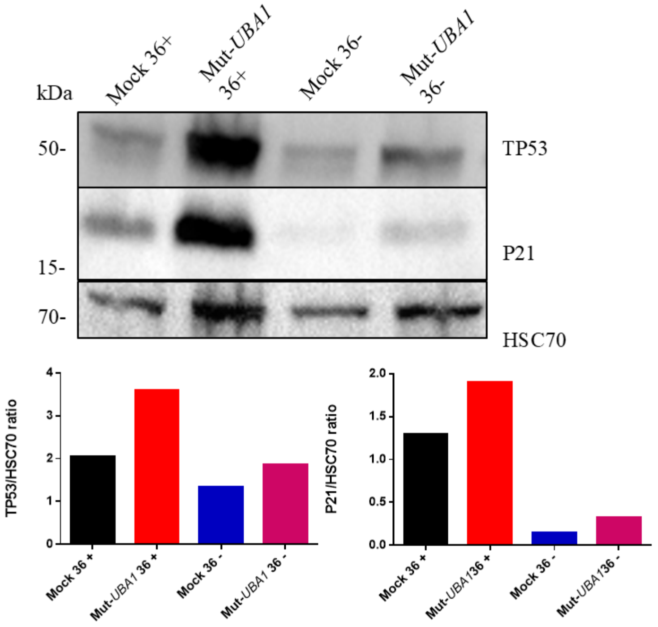


D

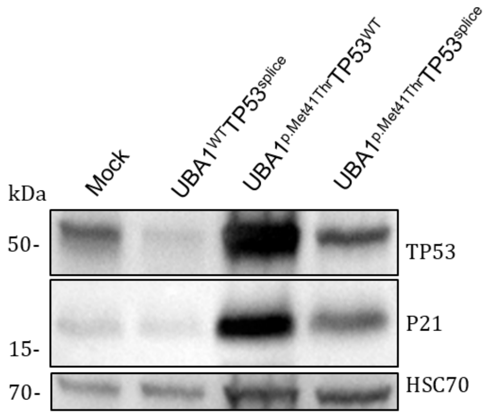

**Supplementary figure 5**: **Exploring the link between TP53 and the death of mut-*UBA1* cord blood erythroid precursors.** A: representative flow cytometry dot blot of annexin V and 7-AAD staining in mock and mut-*UBA1* cord blood $CD36^{pos}$ cells at day 5 of phase II. B: mean aligned sequencing reads counts of *TP53* in RNA-seq of mock and mut-*UBA1* cord blood cells at day 0 of phase II (n = 3 different unrelated donors). C: western blot of TP53 and P21 in sorted $CD36^{pos}$ or $CD36^{neg}$, mock or mut-*UBA1* cord blood cells at day of phase II. D: western blot of TP53 and P21 in the bulk cord blood culture at day 0 of phase II, in mock cells compared to cells solely edited for *TP53* ($UBA1^{WT}$ $TP53^{splice}$), or cells solely edited for *UBA1* ($UBA1^{p.Met41Thr}$ $TP53^{WT}$), or cells simultaneously edited for *TP53* and *UBA1* ($UBA1^{p.Met41Thr}$ $TP53^{splice}$).

#### Supplementary figure 6

A

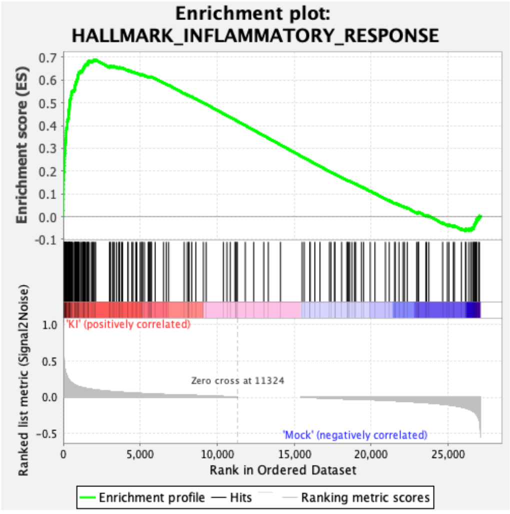


NES 2,59, $p < 10^{-4}$

B

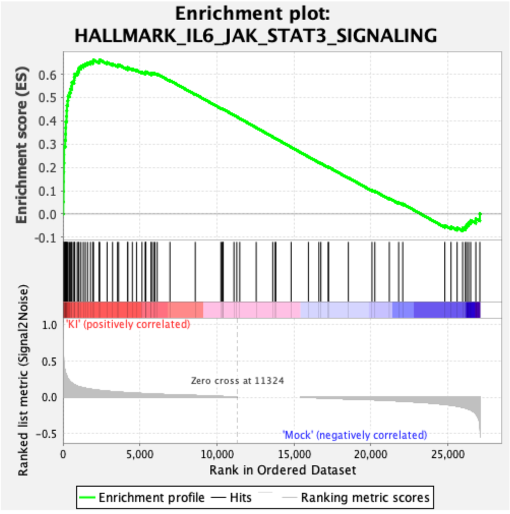


NES 2,21, $p < 10^{-4}$

C

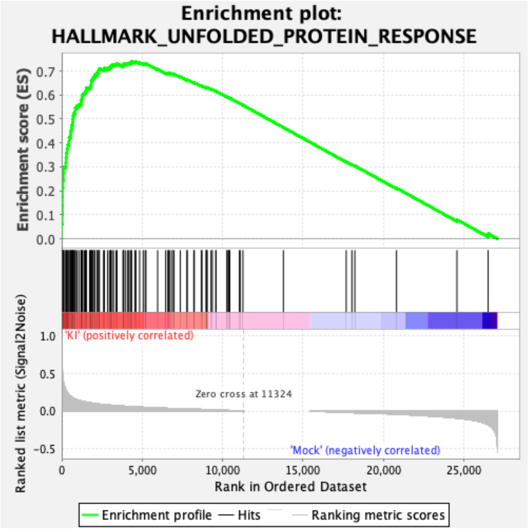


NES 2,57, $p < 10^{-4}$

**Supplementary figure 6: Activation of inflammatory and unfolded protein response (UPR) transcriptional pathways in mut-*UBA1* cord blood cells at day 0 of phase II confirms the relevance of the base-editing model.** A: GSEA of the hallmark inflammatory response in mock and mut-*UBA1* cord blood cells at day 0 of phase II (n = 3 different biological donors). KI = mut-*UBA1*. B: GSEA of the hallmark IL6_JAK_STAT3 in the same conditions.

KI = mut-*UBA1*. C: GSEA of the hallmark unfolded protein response in the same conditions. KI = mut-*UBA1*.

#### Supplementary table 1

| Guide RNAs for *UBA1* base editing | |
|---|---|
| sgRNA1 | 5’ GCCAUUCCCUAGGAAUAGAG 3’ |
| sgRNA2 | 5’ GGCCAUUCCCUAGGAAUAGA 3’ |
| sgRNA3 | 5’ UGGCCAUUCCCUAGGAAUAG 3’ |
| sgRNA4 | 5’ UUGGCCAUUCCCUAGGAAUA 3’ |
| sgRNA5 | 5’ CUUGGCCAUUCCCUAGGAAU 3’ |
| Guide RNA for TP53 base editing | |
| sgRNATP53 | 5’ CCAGACCUCAGGCGGCUCAU 3’ |
| PCR primers for *UBA1* exon 3 sequencing | |
| Forward | 5’ TCCAAAGCCGGGTTCTAACT 3’ |
| Reverse | 5’ TCATGGCCCAACACATACCT 3’ |
| Northern blot probes | |
| 5'ETS$_b$ | 5’-AGACGAGAACGCCTGACACGCACGGCAC-3' |
| 5'ETS-1399 | 5'-CGCTAGAGAAGGCTTTTCTC-3' |
| 5’ITS1 | 5’-CCTCGCCCTCCGGGCTCCGTTAATGATC-3’ |
| ITS1-59 | 5'-CGCGGTGGGGGGGTGGGTGTG-3' |
| ITS1-5.8S | 5'-CTAAGAGTCGTACGAGGTCG-3' |
| 5.8S-ITS2 | 5'-GGGGCGATTGATCGGCAAGCGACGCTC-3' |
| ITS2b | 5’-CTGCGAGGGAACCCCCAGCCGCGCA-3’ |
| ITS2d/e | 5’-GCGCGACGGCGGACGACACCGCGGCGTC-3’ |
| 28S-3'ETS | 5'-CACGCGCGCGCGGACAAACCCTTG-3' |
| 3'ETS | 5'-CTCCCAAACCACGCTCCCCGGACCCCGTCCCGGCCCGGAG-3' |
| 28S | 5’-CCCGTTCCCTTGGCTGTGGTTTCGCTAGATA-3’ |
| 18S | 5’-TTTACTTCCTCTAGATAGTCAAGTTCGACC-3’ |
| 5.8S | 5’-GTTCTTCATCGACGCACGAGC-3’ |

**Supplementary table 1 legend:** list of oligonucleotides (sgRNAs, PCR primers, northern blot probes) used in this study.
